# Interactions between motor cortical forelimb regions and their influence on muscles reorganize across behaviors

**DOI:** 10.1101/2025.01.27.635061

**Authors:** Amy Kristl, Natalie Koh, Mark Agrios, Sajishnu Savya, Zhengyu Ma, Diya Basrai, Sarah Hsu, Andrew Miri

## Abstract

It remains unclear how classical models of motor cortical hierarchy align with emerging evidence of behavioral organization in motor cortex. To address this, we combined optogenetic inactivation, Neuropixels recording, and electromyography to quantify the pattern and influence of activity in the mouse analogs of forelimb premotor and primary motor cortex (RFA and CFA) during reaching and climbing. Results revealed that RFA’s dominant influence on forelimb muscles and on CFA during reaching is replaced by a dominant influence of CFA on muscles and on RFA during climbing, even when forelimb muscle activity during climbing resembles that during reaching. Short-latency influence between regions on excitatory and inhibitory populations in different cortical laminae also showed behavioral specificity. Simultaneous recordings in both areas during climbing revealed a loss of activity timing differences seen during reaching previously interpreted as reflective of hierarchy. These findings demonstrate that hierarchical interactions between forelimb motor cortical regions are behavior-specific.

## INTRODUCTION

Many mammals express a broad array of motor behaviors that vary greatly in the sensory information on which they depend, the muscle activation patterns they involve, and the extent to which they leverage motor programs learned through experience. The flexible adaptation of motor behaviors as context requires further broadens this array. Motor cortex seems to have evolved in size and complexity^1,2^ to improve the performance of diverse behaviors and thereby increase fitness. Yet it remains unclear how motor cortex is organized to achieve this remarkable motor behavioral diversity.

The classical view of a hierarchically organized motor cortex, comprised of “higher-order” premotor (PM) regions that prepare movements executed by “lower-order” primary motor cortex (M1)^3–6^, implies that diverse movements are evoked through a common mode of PM-M1 interaction. This view has an extensive empirical basis that spans anatomy^7–9^, lesion^10–12^, activity^13–16^, and activity perturbation^17,18^. The relative rather than absolute differences revealed by these observations^18–20^ and the recognition of substantial descending projections from PM regions^21–23^ support a more nuanced view of a partial functional hierarchy in which PM regions and M1 interact and each drive downstream motor circuits, but M1 exerts comparatively more influence on movement execution through its denser descending projections. In line with this, we recently reported that the forelimb PM and M1 equivalents in rodents, the rostral and caudal forelimb areas (RFA and CFA) respectively, exhibit asymmetric influence on one another during directional reaching, with RFA having a stronger influence on CFA than vice versa^20^. However, functional interactions between M1 and PM regions have primarily been studied in the context of single-forelimb movements like reaching and grasping. It thus remains unclear whether hierarchical interactions generalize to other behaviors; these sorts of interactions could vary across behaviors, perhaps as an adaptation to promote the efficacy of diverse movements.

Certain existing evidence suggests the possibility that interactions among motor cortical regions and their influence on muscle activity might vary across behavior types. Long-duration stimulation, both electrical^24–26^ and optogenetic^27–29^, in different locations within PM and M1 can evoke movements that resemble different species-typical behaviors, like feeding and climbing. These findings, together with other recent results from inactivation^30–32^ and projection tracing^22,33,34^ in rodents have suggested that PM and M1 regions may be adapted to drive particular behavior types. Our own recent results^35,36^ and those from others^37,38^ have shown that the covariation of M1 neuron firing patterns changes substantially across motor behavior types, which would enable M1’s influence on other regions and on muscles to change. In particular, we have shown that such variation in M1 activity dynamics is observed across a broad range of naturalistic behaviors, including both stereotyped, species-typical behaviors like feeding and grooming, and behaviors that challenge agility and dexterity like climbing and walking across an irregular grid^36^. Other results have suggested that distinct networks oriented toward different behavior types span both frontal motor and parietal cortical regions^39–44^.

Here we addressed whether the hierarchical dynamics observed during directional reaching would generalize to a contrasting ethological behavior. We investigated how mouse RFA and CFA interact and drive muscles in a novel naturalistic climbing context and compared findings to those during directional reaching^20^. To focus on the short-latency influence of endogenous activity patterns in each region, we combined rapid optogenetic inactivation, large-scale multielectrode array (Neuropixels) recording, and electromyography. We found that RFA’s influence on forelimb muscles and on CFA dominated during reaching, while CFA’s influence on forelimb muscles and on RFA dominated during climbing. Across-region influence on excitatory and inhibitory populations in different cortical laminae also changed between behaviors, with only the dominant area during each behavior disproportionately driving inhibitory interneurons in the other area. Simultaneous Neuropixel recordings in both areas during climbing revealed a loss of activity asymmetries seen during reaching and previously interpreted as reflective of hierarchy.

These results demonstrate that interregional interactions in motor cortex change considerably between ethological contexts, perhaps reflective of cortical adaptations that support the diverse motor behavioral repertoire of mammals. In doing so, our results suggest that functional hierarchy is a behavior-specific dynamic, helping to reconcile motor cortical hierarchy with emerging ideas of an ethological organization.

## RESULTS

### Asymmetric influence of endogenous activity on muscles reverses across behaviors

We first addressed whether the relative influence of RFA and CFA on muscle activity varies between behaviors. We briefly inactivated left RFA in climbing mice and compared the immediate effects on contralateral forelimb muscle activity to previous results from inactivating CFA^45^ (Fig. 1a-c). Head-fixed VGAT-ChR2-eYFP mice^46^ (n = 6) climbed at their own pace for water rewards by grabbing and pulling handholds that extend radially from a wheel during daily ∼1 hr sessions (11-20 sessions per animal). Each right handhold’s mediolateral position relative to the mouse is randomized once per wheel rotation, rendering handhold positions unpredictable and thus ensuring adaptive limb movements are needed. An extensive characterization of this behavior is available in a separate paper^45^. To activate Channelrhodopsin-2-expressing (ChR2+) inhibitory interneurons and suppress RFA output^46^, brief blue light pulses (473 nm, 25 ms, 10 mW/mm^2^) were applied to a 2-mm diameter circle on the cortical surface encompassing RFA. Brief inactivation was intended to silence endogenous RFA activity and gauge its immediate influence without substantially disrupting behavioral performance, which could lead to compensatory changes over time. Light pulses were triggered at random times separated by at least 5 seconds during active climbing; these inactivation trials were intermixed with the detection of equivalent events that served as control trials without inactivation. To measure effects of RFA inactivation, muscle activity was simultaneously recorded from chronically-implanted electromyographic (EMG) electrodes in four muscles of the right forelimb (Fig. 1d). We have recently demonstrated that this inactivation method produces remarkably stable effects across trials and sessions^45^.

**Figure 1.**
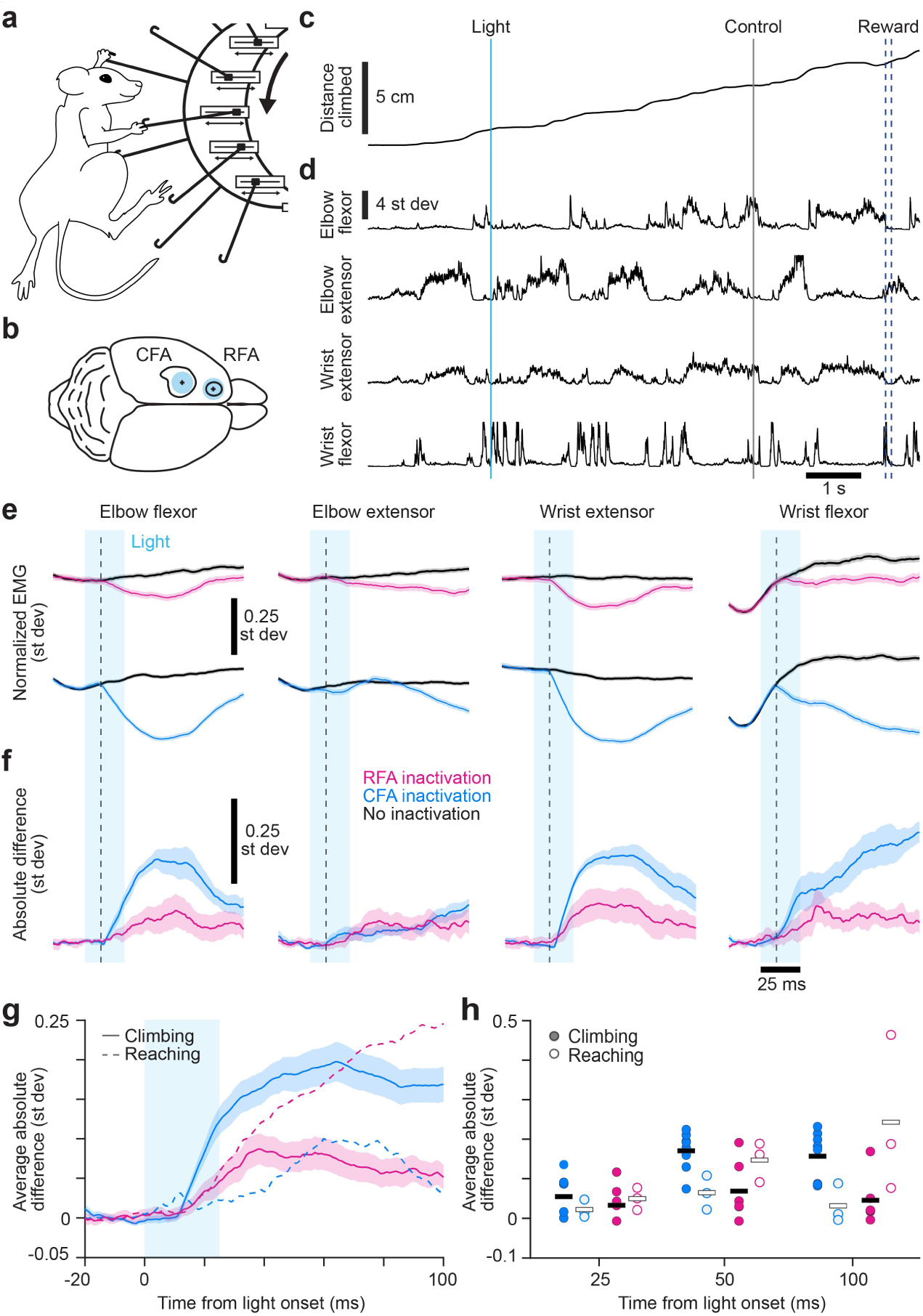
Asymmetry in influence on muscles reverses across behaviors. (**a**) Schematic depicting the climbing behavior. (**b**) Mouse brain schematic depicting the position of the light stimulus on RFA and CFA. (**c**,**d**) Example time series segment of distance climbed with the time of light and control trials, and water reward delivery indicated (**c**) and normalized muscle activity (**d**) over a climbing bout. (**e**) Mean normalized muscle activity (units are st dev) ± SEM across trials without (black) or with inactivation (cyan bar) of RFA (top; n = 7,303 light and 14,289 control trials across 6 animals) or CFA (bottom, n = 9,029 light and 18,397 control trials across 8 animals) for each muscle. The dashed vertical line indicates the shortest latency at which CFA output influences forelimb muscle activity^47^ (10 ms). (**f**) Mean ± SEM absolute difference between inactivation and control trial averages for each muscle for RFA and CFA inactivation (cyan bar) across animals. For baseline correction, the absolute difference was calculated between resampled control trials to estimate the baseline difference expected by chance. (**g**) Mean ± SEM absolute difference between inactivation and control trial averages averaged across all muscles and animals for inactivation (cyan bar) of RFA or CFA during climbing (solid lines). Equivalent means for reaching are overlaid (dashed). (**h**) Average absolute difference across muscles between inactivation and control trial averages at 25, 50, and 100 ms after trial onset for individual animals (circles) and the mean across animals (bars) for climbing (filled circles/black bars) and reaching (open circles/white bars) after inactivation of RFA or CFA.

Averaging normalized muscle activity from thousands of trials across all six mice revealed the effect of RFA inactivation, as compared to our previous results from CFA inactivation (Fig. 1e). Because we have z-scored muscle activity measurements using the mean and standard deviation from each given session, here and throughout we express muscle activity measurements in standard deviations of the recorded signal. For most muscles, averaging randomly timed trials led to flat baselines as the diverse fluctuations in muscle activity across trials averaged out. The wrist flexor’s average had a small deviation at baseline, which may reflect more increases in this muscle’s activity than decreases throughout climbing. As with CFA inactivation^45^, RFA inactivation caused muscle activity to diverge from control levels within 10-20 ms after light onset, similar to the latency of CFA’s fastest influence on forelimb muscles^47^.

In contrast to our previous results during directional reaching for water droplets^20^, we found that during climbing, RFA inactivation had a smaller influence on forelimb muscle activity than CFA inactivation (Fig. 1f,g). The absolute difference in activity between control and inactivation trial averages across animals and muscles was substantially larger for CFA inactivation when measured at 25, 50, and 100 ms after trial onset (Fig. 1g,h; 2.44-fold increase in average absolute difference, p = 0.004 at 25 ms; 2.41-fold increase, p = 0.010 at 50 ms; 3.41-fold increase, p = 0.002 at 100 ms, one-tailed Wilcoxon rank sum test). During reaching, RFA inactivation had a greater influence on muscle activity than CFA (Fig. 1g; 7.38-fold increase, p = 0.002 at 100 ms). The influence of each area on muscles itself varied significantly across behaviors: CFA influenced muscles more during climbing at all time points (p = 0.003 at 25 ms, p = 0.007 at 50 ms, and p = 0.001 at 100 ms), whereas RFA influenced muscles more during reaching than climbing at the latest time point (p = 0.001 at 100 ms). Thus, an asymmetry in RFA and CFA influence on muscle activity seen during reaching reverses during climbing.

We wondered whether these differences were due to the differences in the form of forelimb movements during reaching and climbing. We thus examined whether the influence of each area on muscle activity differed similarly during periods of climbing in which forelimb muscle activity resembled that seen during reaching. In our reaching experiments, inactivation had been triggered by reach onset from rest, though the latency from reach onset to trial onset varied somewhat across trials. To align with this, we identified inactivation and control trials during climbing in which muscle activity in the 50 ms preceding inactivation effect onset was well-correlated with average muscle activity during 50 ms windows beginning at slightly different times during the early phase of reaching in a representative mouse (Supplementary Fig. 1a, see Methods). Averaging muscle activity from these trials aligned on trial onset produced averages akin to those we have previously assembled for reaching (Supplementary Fig. 1b,c). We found that RFA and CFA inactivation during these reach-like trials yielded effects on muscle activity that were highly similar to those seen on other climbing trials (Supplementary Fig. 1d,e). Perhaps not surprisingly then, effects on these reach-like trials were themselves substantially and significantly different from those during reaching onset (CFA inactivation: 4.05-fold increase in average absolute difference, p = 0.004 at 25 ms; 1.50-fold increase, p = 0.019 at 50 ms; 1.69-fold increase, p = 0.003 at 100 ms; RFA inactivation: 0.95-fold decrease, p = 1.76 x 10^-4^ at 100 ms). We repeated this analysis, identifying reach-like trials using muscle activity from two other reaching mice and found similar results (Supplementary Fig. 1f-h). Thus, the differences we observed in inactivation effects between reaching and climbing do not appear to be due to kinematic differences between the behaviors. We also investigated whether variation in inactivation effect size across trials could be explained by differences in the level of muscle activation at trial onset, finding only a weak linear relationship between them (Supplementary Fig. 2).

### Asymmetric interaction between RFA and CFA reverses across behaviors

In our previous work, silencing endogenous activity in RFA during reaching yielded a greater effect on CFA activity than vice versa^20^. To test if this asymmetry in RFA-CFA interaction persists during climbing, we inactivated RFA as above while recording activity across all layers in CFA with Neuropixels 1.0 (13 sessions across 4 mice) or similarly inactivated CFA while recording in RFA (12 sessions across 4 mice). To verify that direct optogenetic inactivation was restricted to the illuminated area, we analyzed spike timing around light onset in narrow-waveform, putative inhibitory interneurons in the recorded area using a modified version of the stimulus-associated spike latency test (SALT, see Methods)^48^ and found no evidence of local optogenetic inactivation in the recorded region (Supplementary Fig. 3). Thus, the suppression observed in the recorded region was assumed to be due to a loss of endogenous input from the inactivated region.

As during reaching, inactivation of each area during climbing caused a strong suppression in the other area’s mean firing rate beginning ∼10 ms after light onset (Fig. 2a-d). Because average firing rates differed between areas and behaviors, we quantified the effect of inactivation by calculating the fractional change in each cell’s average firing rate between inactivation and control trials. We measured this 35 ms after trial onset because inactivation effects began to rebound after this time point (Supplementary Fig. 4). During both reaching and climbing, the majority of neurons in each recorded region showed reductions in firing rate (Fig. 2e).

**Figure 2.**
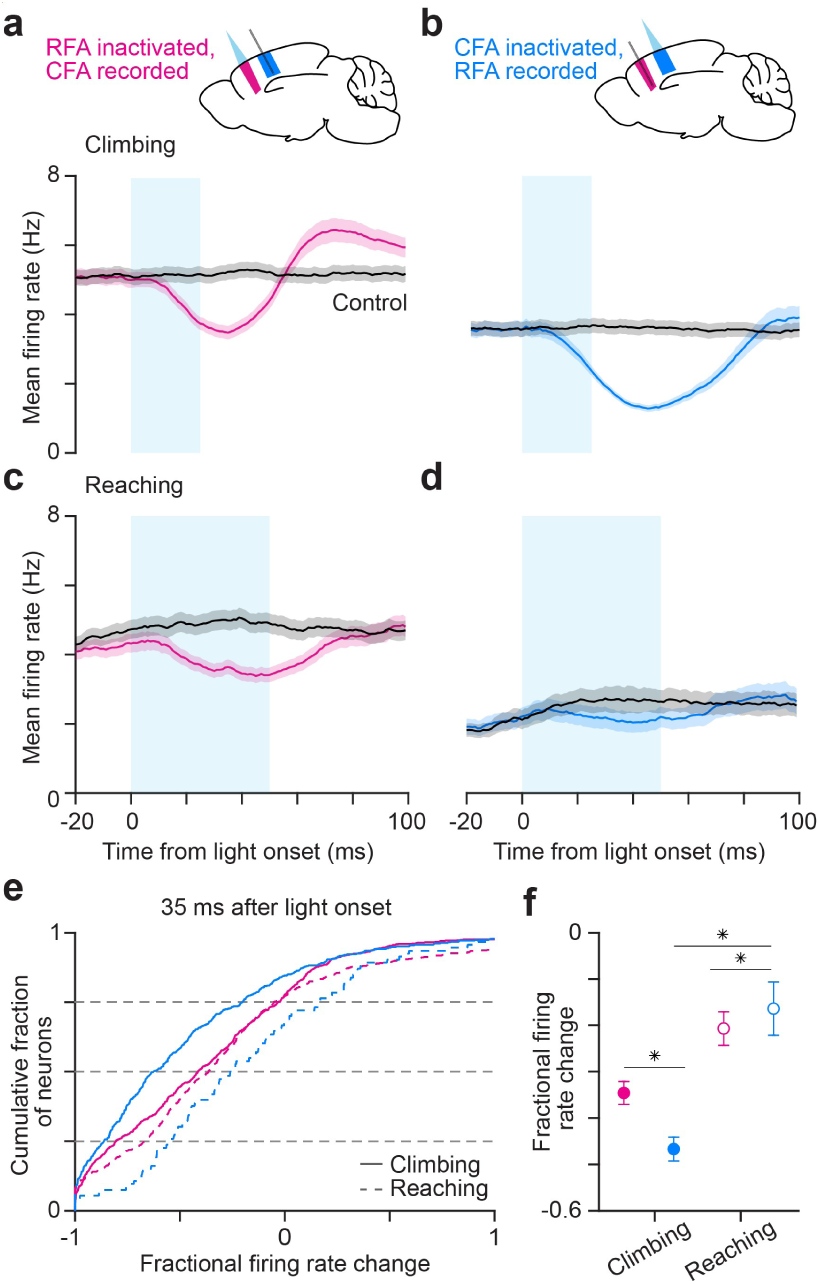
Asymmetry in endogenous neural influence reverses across behaviors. (**a**,**b**) Mean ± SEM firing rate for cells from all four climbing animals after inactivating RFA (cyan bar) and recording in CFA (**a**, 170 cells/animal) or vice versa (**b**, 117 cells/animal) and for control trials. The same number of cells was used from each animal here and in the panels that follow so results equally represent animals despite differing cell yields from recording. (**c,d**) Mean ± SEM firing rate for cells from all three reaching animals after inactivating RFA (cyan bar) and recording in CFA (**a**, 219 cells/animal) or vice versa (**b**, 31 cells/animal) and for control trials. (**e**) Cumulative histograms of the fractional change in firing rate between inactivation and control trials in a 10 second window centered 35 ms after trial onset for neurons in each area after inactivation of the other during climbing or reaching. (**f**) Mean ± SEM fractional change between control and inactivation trials 35 ms after trial onset across neurons in each area after inactivation of the other during climbing and reaching.

Similar to effects on muscles, the asymmetry in RFA-CFA interaction seen in reaching reversed during climbing (Fig. 2f). Whereas CFA inactivation suppressed RFA more than vice versa during climbing (34.97% greater decrease in mean fractional change, p = 3.01 x 10^-6^, Wilcoxon one-tailed rank sum test), RFA inactivation suppressed CFA more than vice versa during reaching (26.22% greater decrease, p = 0.009). This reversal was accompanied by a substantially greater influence of CFA on RFA during climbing than during reaching (184.76% greater decrease, p = 4.17 x 10^-9^). To confirm that the difference in effect size was not due to differing levels of optogenetic inactivation in either area, we recorded from the inactivated region and found similar levels of suppression in RFA and CFA during climbing (Supplementary Fig. 5). Collectively, these results indicate that during climbing, CFA’s influence eclipsed RFA’s to reverse the asymmetry seen during reaching.

### Cell type- and layer-specific interactions between RFA and CFA change across behaviors

Attempts to parse cell type functions in motor cortex have found evidence of task-specific functions: for example, activation of certain inhibitory populations around reach onset^49^. Anatomical studies have revealed that projections between RFA and CFA target a mixture of excitatory pyramidal neurons and inhibitory interneurons^50,51^ in the other area. Asymmetries in the laminar targets of these projections have been interpreted as reflecting hierarchical interactions^8,7,52^, and theoretical considerations suggest the relative engagement of excitatory and inhibitory populations is a key determinant of information flow between regions^53^. We thus took advantage of our large number of recorded cells from all cortical layers to compare inactivation effects on putative pyramidal neurons and inhibitory interneurons, both collectively and in specific layers, between behaviors.

We divided recorded units in each area into wide-waveform, putative pyramidal neurons and narrow-waveform, putative inhibitory neurons based on trough-to-peak spike waveform width using established methods^47,54,55^ (Supplementary Fig. 6). Across animals, 86.1 and 13.9 ± 6.9% of cells were classified as wide-waveform and narrow-waveform, respectively, consistent with histological estimates^56^. Some cells of both types were highly suppressed after inactivation of the other area (Supplementary Fig. 7). Comparing RFA inactivation effects on the two cell types in CFA revealed strong suppression of both cell types during climbing (Fig. 3a,b). Fractional firing rate changes for both types were similar (Fig. 3c,d; wide-waveform: -0.285 ± 0.023; narrow-waveform: -0.333 ± 0.036; p=0.657, Wilcoxon one-tailed rank sum test). However, during reaching, narrow-waveform neurons were more suppressed on average than wide-waveform neurons (138.40% greater decrease in mean fractional change, p = 0.035). Thus, another prominent asymmetry during reaching was not seen during climbing.

**Figure 3.**
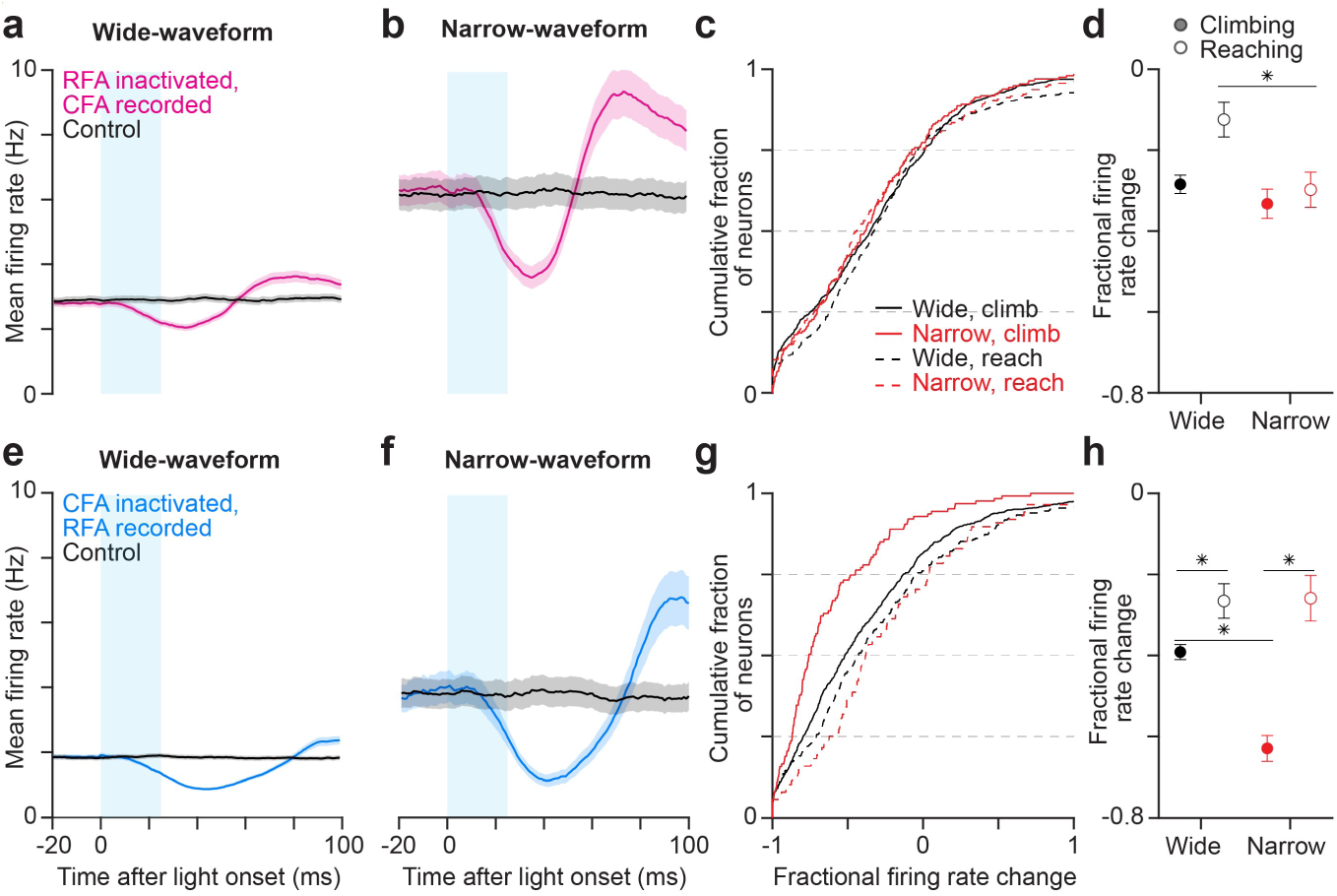
Only the dominant area exerts a larger effect on inhibitory interneurons. (**a**,**b**) Mean ± SEM firing rate for all wide-(**a**, n = 1299) or narrow-waveform (**b**, n = 230) neurons in CFA after RFA inactivation and for control trials. (**c**) Cumulative histogram of the fractional change between inactivation and control trials 35 ms after trial onset for wide-(n = 850) and narrow-waveform (n = 194) neurons in CFA after RFA inactivation during climbing or reaching. (**d**) Mean ± SEM fractional firing rate change between inactivation and control trials 35 ms after trial onset for all wide- and narrow-waveform neurons in CFA after RFA inactivation during climbing and reaching (n = 637 wide, n = 206 narrow). (**e**,**f**) Same as (**a**,**b**), but for RFA neurons (n = 1583 wide, 187 narrow) recorded during CFA inactivation. (**g**,**h**) Same as (**c**,**d**), but for RFA neurons (n = 904 wide, 127 narrow during climbing; n = 316 wide, 74 narrow during reaching) during CFA inactivation.

The effects of CFA inactivation on both cell types in RFA during climbing (Fig. 3e,f) were also very different from those seen during reaching (Fig. 3g). On average, both types were much more strongly suppressed during climbing (Fig. 3h; wide-waveform: 47.37% greater decrease in mean fractional change, p = 0.006; narrow-waveform: 143.24% greater decrease, p = 6.57 x 10^-9^). However, here the effect on narrow-waveform neurons was substantially and significantly higher than the effect on wide-waveform neurons during climbing (60.71% greater decrease, p = 2.15 x 10^-7^) and not reaching. Thus, whereas cell type-specific effects of RFA on CFA were seen only during reaching, the relatively strong, cell type-specific effects of CFA on RFA were seen only during climbing. These type-specific effects have a common form: during each behavior, the dominant area (CFA during climbing, RFA during reaching) exerts a larger effect on inhibitory interneurons in the other area, while the non-dominant area exerts a balanced effect across cell types. The fact that silencing either area still has a net suppressive effect on the other area (Fig. 2) indicates that any disinhibition caused by loss of drive to interneurons is overcome by the loss of drive to the approximately six-fold larger population of excitatory neurons.

Dense projections from RFA to CFA’s layer 5 that outnumber CFA’s projections to RFA’s layer 5 are thought to reflect a stronger influence of RFA on descending output from CFA than vice versa^7,8,52^. We thus examined the influence of each area on narrow- and wide-waveform neurons assigned to layers 2/3 (L2/3), 5 (L5), and 6 (L6), using conservative depth criteria for assignment to each layer (Supplementary Fig. 8). An ANOVA indicated a significant dependence of effects on layer for inactivation of each region (p=0.001 for RFA inactivation, p = 2.069 x 10^-12^ for CFA inactivation). Based on this, we further examined how effects of inactivating each area varied across cell groups defined by layer and waveform type.

RFA inactivation during climbing had broad-based effects across laminar groups in CFA, as all groups had mean fractional change values between -0.248 and -0.483 (Fig. 4a-c, Supplementary Table 1). Narrow-waveform neurons in L5 were significantly more suppressed than narrow-waveform neurons in other layers, while effects were instead similar for wide-waveform neurons across layers. To compare across behaviors, we calculated the mean difference in fractional firing rate change between behaviors for each laminar group (Fig. 4d; Supplementary Fig. 9). Wide-waveform neurons in L2/3 and L5 and narrow-waveform neurons in L2/3 showed significantly larger drops in firing rate, indicating that RFA had a greater influence on these groups during climbing than reaching.

**Figure 4.**
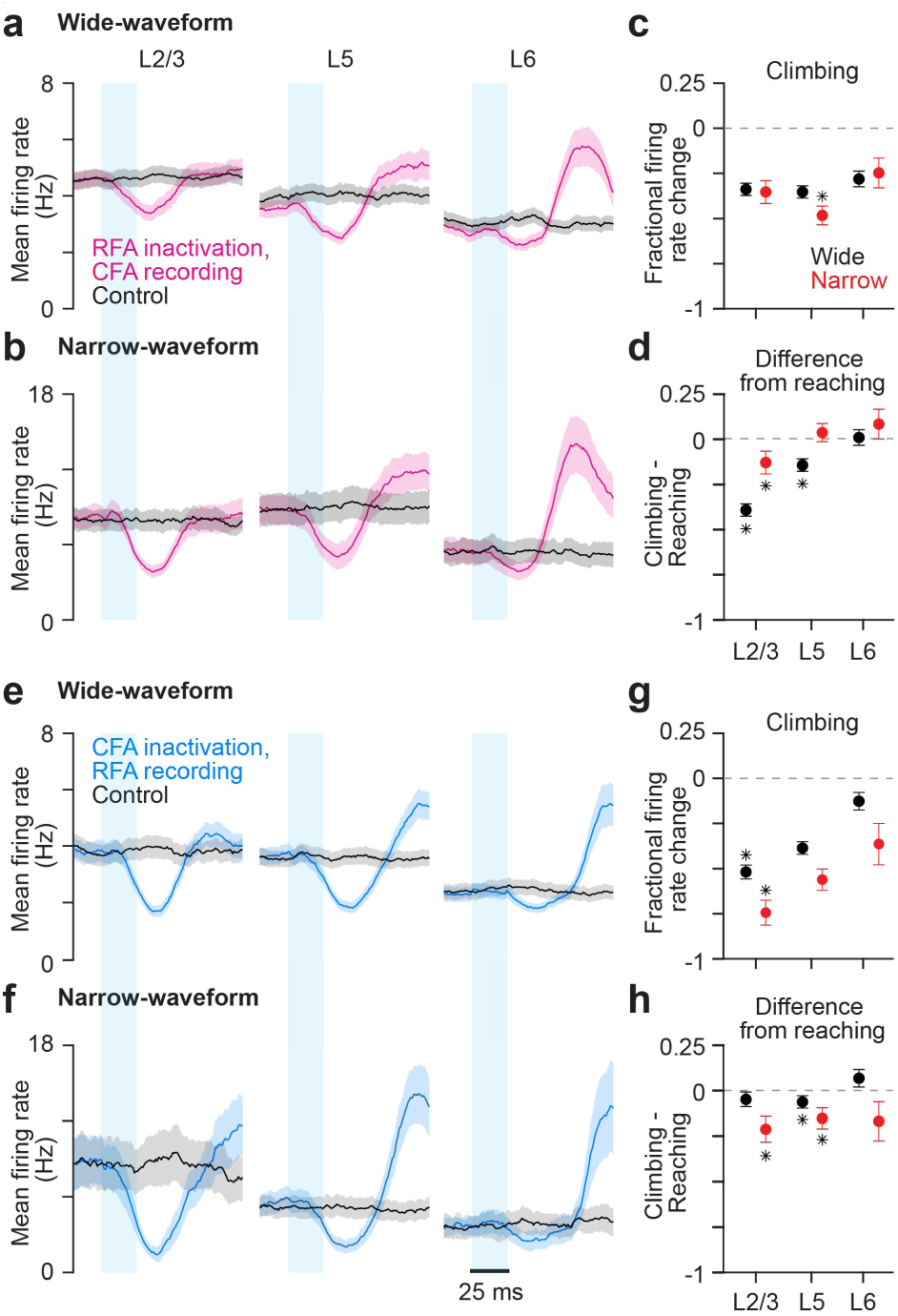
Effects of inactivation on cell types vary by cortical layer. (**a**,**b**) Mean ± SEM firing rate for wide- (**a**; 324 in L2/3, 167 in L5, 130 in L6) and narrow-waveform (**b**; 60 in L2/3, 52 in L5, 47 in L6) neurons in CFA assigned to each laminar cell group after RFA inactivation, and for control trials. (**c**) Mean ± SEM fractional firing rate change between inactivation and control trials 35 ms after trial onset for all neurons in different laminar cell groups in CFA after RFA inactivation during climbing. (**d**) Mean ± SEM difference between the fractional firing rate change of neurons in each laminar cell group 35 ms after trial onset and the mean fractional change of the equivalent group during reaching. (**e**)-(**h**) Same as (**a**)-(**d**), but for RFA neurons (wide: 103 in L2/3, 156 in L5, 132 in L6; narrow: 12 in L2/3, 34 in L5, 12 in L6) recorded during CFA inactivation, instead of vice versa.

Examining the effects of CFA inactivation on each laminar cell group in RFA, we found that effects were again broad-based (Fig. 4e-g). In this case we saw a clear trend of stronger effects in more superficial layers (Fig. 4g). Both wide- and narrow-waveform neurons in L2/3 were significantly more suppressed than all other neurons of their respective waveform types (Fig. 4g). The effect on narrow-waveform neurons in L2/3 was particularly strong, with a mean fractional change of -0.741. Inactivation effects were larger during climbing than reaching in narrow-waveform cells in L2/3 and L5 and wide-waveform cells in L5 (Fig. 4h). Thus, interregional influence varies across layers, and the change in this influence between behaviors varies substantially across layers. Importantly, CFA’s effects on L5 in RFA were comparable to RFA’s effects on CFA L5. This is inconsistent with the view that RFA retains a stronger influence on descending output from CFA during climbing.

### Asymmetries in RFA and CFA activity change across behaviors

Several asymmetries between activity in PM and M1 have previously been interpreted as potentially reflective of motor cortical hierarchy, including those seen during single-forelimb motor behaviors in rodents^13,57–60^. Indeed, we also found asymmetries of this sort between activity in RFA and CFA during reaching^20^. We thus examined whether such asymmetries might differ during climbing.

Using Neuropixels, we simultaneously recorded activity in RFA and CFA together with forelimb muscle activity during climbing (16 sessions from 10 wild-type mice). Basic firing features resembled those seen during reaching, as most neurons in both areas fired more frequently during active climbing than not (Fig. 5a,b), and mean firing rates were broadly distributed but higher on average in narrow-waveform neurons (Fig. 5c). During reaching, a larger fraction of neurons in RFA than in CFA had firing that was significantly correlated with the activity of at least one contralateral forelimb muscle. However, during climbing, there were more neurons in CFA meeting this criterion (Fig. 5d; p=0.012, Wilcoxon one-tailed rank sum test). We also found that the difference in the fractions significantly correlated in RFA and CFA was itself significantly different between behaviors (Fig. 5e; climbing +0.069, reaching -0.092, p=0.008).

**Figure 5.**
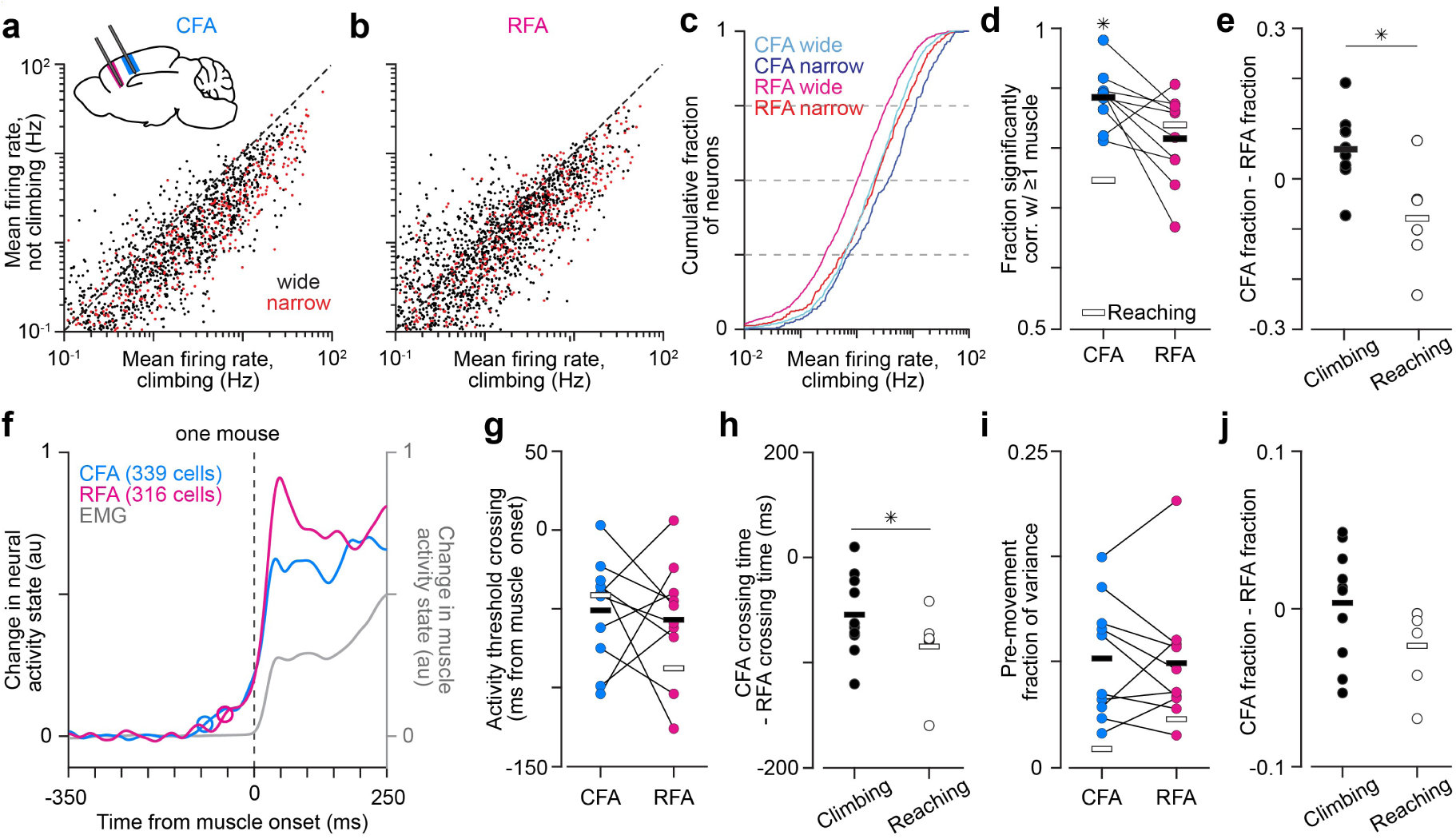
Relations between firing in RFA and CFA change across behaviors. (**a**,**b**) Scatter plots of the firing rates for cells recorded in CFA (**a**) and RFA (**b**) during periods of active climbing versus all other time periods for wide- and narrow-waveform neurons. (**c**) Cumulative histogram of the mean firing rates of wide- and narrow-waveform neurons recorded in CFA and RFA during climbing. (**d**) The fraction of neurons recorded in CFA and RFA for each animal during climbing (circles) and the mean across animals (black bars) whose firing rate time series was significantly correlated with that of at least one muscle (p-value threshold < 0.05). The mean fraction in each area across animals during reaching is overlaid (white bar). (**e**) The difference between the fraction of neurons in CFA and RFA whose activity was significantly correlated with that of at least one muscle for each animal (circles) and the mean across animals (bars) during climbing (n = 10 mice) and reaching (n = 6). (**f**) For an example mouse, the normalized activity change from baseline summed across the top three principal components (PCs) for all recorded CFA or RFA neurons, and the top PC for muscle activity surrounding muscle activity onset. Circles indicate the time of estimated activity onset in each area. (**g**) Time from muscle activity onset at which the neural activity change from baseline exceeded a low threshold for each animal (circles) and the mean onset time across animals (bars). The mean onset times across animals during reaching is overlaid (white). (**h**) The difference between activity onset times in CFA and RFA for each animal (circles) and the mean across animals (bars) during climbing and reaching. (**i**) The activity variance in the 150 ms before muscle activity onset, defined as a fraction of the total activity variance from 150 ms before to 150 ms after muscle activity onset for each animal (circles) and the mean across animals (bars). (**j**) The difference between the fraction of variance captured before muscle activity onset in CFA and RFA for each animal (circles) and the mean difference across animals (bars) during climbing and reaching.

We next compared RFA and CFA activity preceding movement onset across behaviors. As we had previously done for reaching, we quantified the activity change around muscle activity onset in each region, using the absolute change from baseline in trial-averaged neural activity summed over the top 3 principal components (PCs, Fig. 5f). Unlike during reaching, when RFA activity rose significantly earlier than CFA activity across animals, during climbing there was no clear and consistent leader at onset (Fig. 5g); RFA led CFA in some animals while CFA led RFA in others. The same was true regardless of the thresholds used for onset detection, when using mean firing rates instead of PCs for onset detection, and when looking specifically at wide- and narrow-waveform neurons (Supplementary Fig. 10). The difference in onset time between CFA and RFA was near zero on average during climbing, and significantly different from reaching (Fig. 5h; climbing +1.0 ms, reaching -46.4 ms, p = 0.044). This indicates that the lag between activity changes in RFA and CFA preceding reaching onset does not generalize to climbing.

We next examined the relative fraction of activity variance that occurs before climbing onset in each area. With reaching, we had previously found a significantly higher fraction in RFA, in line with previous observations of more pre-movement activity in PM regions^57,58,60,61^. From the summed, baseline-subtracted time course of the top three PCs (Fig. 5f), we computed the area under the curve in the 150 ms prior to movement onset and compared it to the area under the curve from 150 ms prior to 150 ms after onset. In contrast to reaching, and despite a larger number of observations (10 mice vs. 6 during reaching), the relative activity variance before climbing onset was not significantly different between regions (Fig. 5I; p = 0.661), and the difference between regions was close to zero (Fig. 5j; difference = 0.004 ± 0.011).

Lastly, as we had previously done for simultaneous RFA and CFA recordings during reaching, we used delayed latents across groups (DLAG)^62^ to decompose activity in each region into latent variables (components) that are either unique to activity in one region (within-region) or shared between regions at an arbitrary temporal delay (across-region), and then probed for asymmetry between the across-region components that appear earlier in each region. We decomposed each area’s activity (26 climbing sessions from 14 mice) into four across-region components (negative delays defined as RFA-leading) and four within-region components. Fig. 6a shows results for across-region components identified from recordings in one mouse (two sessions). Across-region components whose delay was not significantly different from zero according to a permutation-based statistical test^62^ (open circles) and components with an absolute delay greater than 60 ms were excluded from further analysis to focus on fast-timescale interactions; 60 ms had been chosen based on the average onset delay between RFA and CFA during reaching^20^. As during reaching, DLAG models across animals found across-region components in which either direction lead (Fig. 6b), consistent with bidirectional interaction between RFA and CFA.

**Figure 6.**
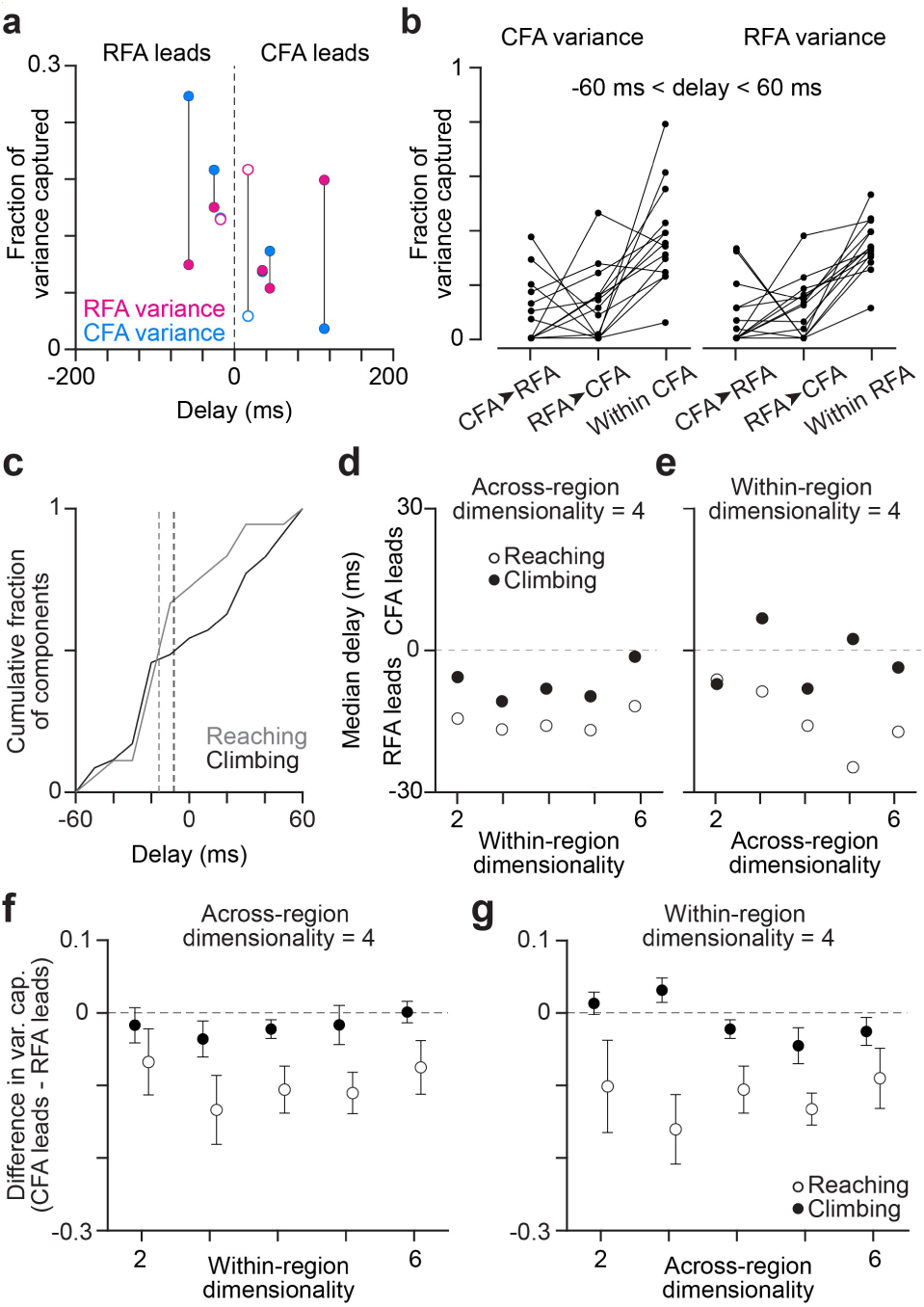
Variance captured by activity patterns shared at a lag changes across behaviors. (**a**) Scatter plot illustrating the delays of all across-region components detected by DLAG versus the fractional variance captured in each region for one mouse (2 sessions). Open circles indicate that the time delay was not significantly different from zero. Lines connect variance capture for individual components. (**b**) Fractional variance capture of CFA activity (left) and RFA activity (right) by across-region components in which CFA activity led RFA (CFA➤RFA), or RFA led CFA (RFA➤CFA), and by within-region components, in DLAG models fit with four across-area and four within-area components. Connected dots indicate individual mice (n = 14). (**c**) Cumulative histogram of all significant component delays from all mice during climbing and reaching from DLAG models fit with four across-area and four within-area dimensions. Dashed lines indicate the median delay for each behavior. (**d**,**e**) The median delay across all significant component delays when varying within- (**d**) or across-region (**e**) dimensionality during reaching (open circles) and climbing (filled circles) with the other number of dimensions set to four. Positive values indicate CFA led more components, whereas negative values indicate RFA led more components. (**f**,**g**) Mean ± SEM difference between the variance captured in RFA activity by CFA-leading components and the variance captured in CFA activity by RFA-leading components across animals, for reaching (open circles) and climbing (filled circles) when varying the within-region (**f**) or the across-region (**g**) dimensionality. Values below zero indicate RFA-leading components captured more variance in CFA activity than vice versa.

Among these across-region components, we probed for asymmetries in their temporal delays and downstream variance capture like those seen during reaching. We first asked whether more across-region components originated in one area or the other by analyzing the distribution of component delays (Fig. 5c). During reaching, there were more RFA-leading components than CFA-leading components, evidenced by a skewed cumulative distribution of delays with a median of -15.9 ms that was significantly lower than zero^20^. However during climbing, the distribution of delays had a median of -8.09 ms, which was not significantly different from zero (p = 0.987, two-tailed Wilcoxon signed rank test). This finding was consistent across models fit with different numbers of within- and across-region components (Fig. 6d,e, Supplementary Table 1). Therefore, the asymmetry in across-region component origin observed during reaching was reduced.

Finally, we compared the variance captured in the lagging region by CFA-leading and RFA-leading components because RFA-leading components had captured more variance during reaching. To compare the asymmetry in each region’s downstream variance capture between behaviors, we calculated the difference between variance captured in the lagging region by CFA- and RFA-leading components for each animal and found a significant decrease in these differences during climbing compared to reaching (p=0.005, one-tailed Wilcoxon rank sum test). This finding was also generally consistent across models fit with different numbers of within- and across-region components (Fig. 6f,g; Supplementary Table 1). Thus in general, the asymmetries in activity timing seen during reaching were diminished or absent during climbing. However, in contrast with results from optogenetic inactivation, these asymmetries did not reverse direction. This divergence between observations from activity perturbation and activity timing analysis agrees with our previous findings during reaching^20^.

## DISCUSSION

We have assessed hierarchical neural dynamics through the fast timescale influence of endogenous activity patterns in forelimb motor cortex during two contrasting motor behaviors. Comparing optogenetic inactivation of RFA and CFA during reaching and climbing revealed that the asymmetry in functional influence between the regions reverses direction between behaviors. RFA’s influence on muscles and on CFA dominated during reaching, while CFA’s influence on muscles and on RFA dominated during climbing. This finding held even when considering only climbing movements that resembled reaching. Additionally, the dominant area during each behavior exerted a larger effect in the other area on putative inhibitory interneurons than on putative pyramidal neurons, in contrast to the non-dominant area. Finally, analysis of concurrent neural activity in both areas during climbing revealed that asymmetries seen during reaching were not apparent during climbing. Our results demonstrate that interregional interactions can change substantially between ethological contexts, indicating that motor cortical hierarchy may itself be a behavior-specific dynamic.

That motor cortical hierarchy may be behavior-specific aligns with the idea of “emergent hierarchies” within motor cortex originally proposed to explain how different regions driving different motor behavior types interact during complex behaviors that involve multiple types^63^. In this view, a subregion of motor cortex that commands a distinct type might need to recruit other subregions of motor cortex to drive specific subcomponents of the full behavior. For example, if CFA controls climbing and RFA reaching, CFA activation may recruit RFA during reaches for upcoming handholds. Consistent with this, RFA’s influence on CFA and muscles was similar across behaviors, in contrast to CFA’s influence on RFA and muscles. This sort of behavior-specific across-region recruitment would enable the engagement of the particular downstream motor circuits uniquely engaged by projections from a given cortical region. As such, it may reflect a tradeoff between the efficiency gains from the proximity of neurons interacting during particular behaviors and the costs of downstream projections that would redundantly engage the same motor effector circuits.

Behavior-specific hierarchy can account for several previous findings that have suggested functional distinctions between RFA and CFA. In mice, stimulation for hundreds of ms in CFA evokes rhythmic oscillatory movements of the contralateral forelimb that resemble locomotion, while such stimulation in RFA evokes reach- and grasp-like movements^28,29,64^. In rats, rhythmic movements have not been observed from CFA stimulation^27,30^, but separate CFA regions are oriented toward forelimb elevation, advance, and retraction, which together could produce locomotor movements^30^. Activity measurements have shown a preferential activation of neurons in RFA during reach and grasp movements, while a much larger fraction of neurons in CFA are not strongly modulated during these movements^31^. Though doubts have been raised regarding the interpretation of long-duration stimulation effects with respect to motor cortical function^65^, our results here examining the short-latency influence of endogenous activity during behavior, perhaps surprisingly, validate such interpretations. Interestingly, behavior-specific interregional dynamics in motor cortex would seem to contrast with emerging ideas of visual cortical organization, which emphasize “task-general” sensory representation as a significant organizational constraint^66^. As we note above though, we expect that the behavioral specialization in motor cortex exists to some extent in tension with other adaptations.

How might a region’s relative influence on different cell types and laminae in another region, or on downstream motor effector circuits, change across behaviors? One possibility is that changes in the covariation of activity patterns output from a region could enable differential effects in target regions. Such changes have previously been proposed as a mechanism for selectively engaging effector circuits during movement but not movement preparation^67^, during some motor behaviors and not others^47^, and to drive different activation patterns in muscle groups^35^. Moreover, there is now a broad range of results across movement contexts describing such changes in motor cortical activity covariation, especially across behaviors that differ substantially in motor output pattern^35,68–71^. Similar changes have also been described in regions beyond motor cortex^72–75^. Such a role for covariation changes is consistent with the idea that brain regions communicate through discrete activity subspaces^76^. Here these subspaces would be engaged in a behavior-specific manner to differentially influence different cell types and lamina in target regions.

One apparent discrepancy remains between results from optogenetic inactivation, which revealed a hierarchical relationship between areas that reversed direction across behaviors, and from activity timing analysis, which revealed asymmetric interactions during reaching but more symmetric interactions during climbing. One potential explanation is that the earlier activity changes are not prerequisite for a feedforward influence; activity timing differences could instead reflect differential amounts of preparatory activity unrelated to drive to the other area^63^. Our DLAG results are more puzzling because although the strong asymmetry from reaching was reduced in climbing, it remained slightly biased toward a larger RFA-to-CFA drive. This could reflect the fact that DLAG only detects activity patterns having the same form in both regions, which may not hold for much of the RFA activity driven by CFA output.

There are several prominent differences between reaching and climbing that may inform motor cortical organization and the changes in interregional dynamics we observed. One key difference is the sensory requirement of each behavior. Climbing in our paradigm is adaptive and substantially non-stereotyped, depending heavily on somatosensory signals. The randomized positions of the right handholds are detectable by the whiskers and hands, and mice frequently re-adjust their grip on the handholds after the initial grasp. Past studies suggest that CFA is specifically involved in adaptive or corrective movements that require such tactile feedback^77,78^. CFA is also highly connected with adjacent forelimb primary somatosensory cortex (S1)^79^. The reaching paradigm we have used, on the other hand, involves a pre-planned, largely stereotyped movement that depends on a preceding visual cue dictating where to reach and an auditory cue dictating when to reach. Secondary motor cortex, which contains RFA, is coupled with visual, auditory, and prefrontal cortices^6,80–83^, positioning RFA to integrate their input to plan and execute a targeted reach. A second key difference is the time course of the behavior: directional reaching is a largely ballistic movement whereas climbing is a continuous, rhythmic behavior. Parcellation of ballistic and rhythmic movements is also seen in vibrissal cortex, where stimulation of whisker representations in M1 and S1 evokes rhythmic whisking and prolonged whisker retraction, respectively^84^. CFA might preferentially engage downstream pattern-generating circuits to enable adaptive control of rhythmic behavior. Lastly, limb involvement also differs substantially: reaching involves extending a single limb whereas climbing involves coordinating all four limbs together. CFA is adjacent to cortical representations of the hindlimbs and whiskers^85^, and their proximity could enable efficient communication between regions to coordinate climbing behavior.

The behavioral organization we demonstrate here challenges a broad consensus^8,86–92^ that RFA is the rodent analog of the supplementary motor area (SMA) in primate cortex. SMA in primates is thought to be more involved in self-initiated actions^93,94^, bimanual coordination^95,96^, and postural stability during locomotion^97,98^. All of these features are prominent in our climbing paradigm but not our reaching paradigm. Moreover, as we mention above, our results substantiate a preferential role for RFA in reach and grasp movements, which aligns more with the function of other premotor regions but not SMA in primates^99–101^. Results from neural activity measurement in^102^ and tracing projections from^9^ RFA have revealed parallels to primate dorsal and ventral PM regions more so than SMA. Bimanual effects of electrical stimulation in CFA have also been described^42^. We thus propose that RFA is more appropriately viewed as a homolog of primate dorsal and/or ventral PM regions, while CFA aligns functionally more so with SMA.

Our results help clarify what we describe as (at least) a tripartite behavioral map in rodent motor cortex. In addition to regions adapted for locomotion (CFA) and reach-to-grasp movements (RFA), recent results are consistent with the existence of a third, rostrolateral region that preferentially controls oromanual eating^33,64,103^. The positioning of this eating region adjacent to RFA could facilitate interactions between eating and grasping circuits. This adjacency is itself reminiscent of primate motor cortical organization, where regions hypothesized to control eating^104^ and hand-to-mouth movements^99^ are close to one another. However, a tripartite behavioral map may only reflect the primary, or “highest,” level of behavioral organization. In line with this, some evidence suggests a topographic distinction between subregions in RFA that control reaching and grasping^27,31^, in addition to the proposed functionally distinct subregions of CFA referenced above.

The timescale on which functional hierarchy changes also remains to be seen – changes may occur on fast timescales as task demands vary. In fact, recent results from modeling activity in primate PM and M1 suggest that hierarchical interactions may change between the preparation and execution of a reach^105^, and modeling of activity in V1 and V2 has suggested similar changes in hierarchical interactions between these regions on fast timescales^62^. We expect understanding of ethological adaptation at finer spatial and temporal granularity to emerge as techniques for more focal activity perturbation continue to develop.

## ACKNOWLEDGEMENTS

A.K. was supported by the Dr. John N. Nicholson Fellowship and NIH training grant 2T32MH067564. A.M. was supported by a Searle Scholar Award, a Sloan Research Fellowship, a Whitehall Research Grant Award, The Chicago Biomedical Consortium with support from the Searle Funds at The Chicago Community Trust, the Simons Foundation, and NIH grant DP2 NS120847.

## AUTHOR CONTRIBUTIONS

A.K. and A.M. devised the project and designed experiments. A.K., N.K., M.A., S.S., Z.M., and D.B. performed experiments. A.K, Z.M., and S.H. analyzed data. A.K. and A.M. wrote the manuscript.

## DATA AVAILABILITY

The data that support the findings of this study will be uploaded to an open data archive like DANDI upon publication.

## CODE AVAILABILITY

All Matlab code used for data analyses will be made available on the Miri lab’s GitHub page upon publication.

## STAR METHODS

### Experimental Animals

All experiments and procedures involving animals were performed in accordance with NIH guidelines and approved by the Institutional Animal Care and Use Committee of Northwestern University. A total of 71 adult male mice were used in reported experiments and early pilot studies to establish methodology. Strain details and number of animals in each group are as follows: 25 VGAT-ChR2-EYFP line 8 mice (B6.Cg-Tg(Slc32a1-COP4*H134R/EYFP) 8Gfng/J; Jackson Laboratories stock # 014548) and 46 C57BL/6J mice (Jackson Laboratories stock #000664). All mice were individually housed under a 12 hr light/dark cycle during the course of the experiment. At the time of reported measurements, animals were 12-24 weeks old and weighed approximately 24**-**30 g. All animals were used for the first time in this experiment and were not previously exposed to pharmacological substances or altered diets.

### Climbing Apparatus

The apparatus was located inside a sound attenuating chamber (H10-24A, Coulbourn). Experimental control was performed using the Matlab Data Acquisition Toolbox, a NI PCI-e-6323 DAQ, and two Arduino Duos. The climbing apparatus consisted of a 3D-printed cylindrical wheel with 36 left-right alternating handholds positioned 12 degrees apart from each other. The handholds on the animal’s left were affixed to the wheel and the handholds on the animal’s right were attached to linear actuators (L-12-30-50-12-I, Actuionix). One end of the wheel was attached to a shaft angular encoder (A2-A-B-D-M-D, U.S. Digital) which measured angular position signals. These signals were delivered to the Arduinos to track the location of each right handhold. When each right handhold reached a radial position 180° away from the mouse, its linear actuator moved the handhold to a random position along the wheel’s width (mediolaterally for the mouse). The other end of the wheel was supported by a slip ring (SR20M-L T, Michigan Scientific) that facilitated the transfer of signals to and from the actuators.

Water rewards were dispensed with a solenoid valve (161T012, NResearch) attached with plastic tubing to a lick tube (01-290-12, Fisher). The dispensation was triggered by Matlab through the NI PCI-e-6323 DAQ. Water dispensation was calibrated to ensure that each water reward contained 3 μl of water. During behavior, mice were head-fixed in an upright position facing the wheel so that all four limbs could easily grab onto the handholds, and so handholds would brush past the mouse’s whiskers during climbing. The lick tube was positioned under the mouse’s mouth to allow water reward delivery during behavior.

### Headplate Implantation

Under anesthesia induced with isoflurane (1-3%), mice had 3D printed plastic head plates^106^ affixed to their skull using dental cement (Metabond, Parkell). During the implantation surgery, marks were made on one side of the headplate at a known distance from bregma to facilitate the positioning of craniotomies for inactivation or recording made in a later surgery. A subset of mice also had EMG electrodes implanted during headplate implantation surgery (see below). After recovery from surgery, mice began behavior training.

### Behavior Training

Mice were first placed on a water schedule in which they received 1 ml of water per day after recovery from headplate implantation. At least 4 days after the start of the water schedule, mice were acclimated to handling by the experimenter using established procedures^107^. Over the next two days, mice were acclimated to head-fixation via daily sessions in which they were head-fixed and restrained in a hutch positioned directly in front of the climbing wheel and provided water rewards at regular intervals.

After this acclimation period, mice had daily training sessions on the climbing wheel. Training was broken into 3 stages. The first stage (1-3 sessions) involved mounting the animal on the wheel and manually triggering rewards with a key-press whenever the mouse made any sort of climbing movement, to encourage mice to climb on the wheel and associate climbing with water rewards. Most mice began climbing within the first session, however some mice required additional sessions in this stage. Once a mouse was able to sustain regular climbing bouts, it progressed to the second stage. During this stage of training, the right handholds were set in a fixed mediolateral position, and mice were encouraged to climb on the wheel for increasingly long periods of time by a Matlab script that triggered water reward dispensation after bouts of continuous climbing longer than a distance threshold. The distance threshold adapted as follows over the course of a session: First, it was set to a low value, and the mouse received 1 water reward each time it met that threshold. After the mouse met that threshold 10 times, the threshold was modified based on the distance climbed in the previous 10 climbing bouts. If the distance was greater than the 25th, 60th, or 90th percentile value for the previous 10 bouts, the mouse received 1, 2, or 4 rewards, respectively; otherwise, the mouse received no water reward. Mice were transitioned to the next stage after two days of climbing with fixed handholds. During the final training stage, the right handhold positions were set to randomly reset their mediolateral position after rotating past the mouse and the same reward scheme was used as in the second stage.

### EMG Recording

EMG electrodes were hand-fabricated for forelimb muscle recording using established procedures^47,108,109^. Each set consisted of four pairs of electrodes, each fabricated as follows. Two 0.001” braided steel wires (793200, A-M systems) were knotted together. The length of wire between the knot and connector was 3.5 cm for elbow flexor and extensor muscles and 4.5 cm for wrist flexor and extensor muscles. On one wire in each pair, insulation was stripped 1 to 1.5 mm away from the knot; on the other wire, insulation was stripped 2 to 2.5 mm away from the knot, yielding two non-overlapping segments of stripped wire that would ultimately be implanted in muscle tissue. The ends of the wires on the opposite side of the knot were soldered to an 8-pin miniature connector (33AC2364, Newark). The other ends had 0.5 cm of insulation stripped from their ends and were then twisted and crimped inside a 27-gauge needle to facilitate implantation. Mice were implanted with EMG electrodes and headplates in the same surgery. The connector was cemented to the posterior edge of the headplate to allow access during behavior experiments, and the electrode wires were led under the skin from the incision on the animal’s neck to skin incisions on the forelimb. The needle on each electrode was used to insert the electrode proximal-to-distal into the target muscle. Insertions targeted the biceps (elbow flexor), triceps (elbow extensor), extensor carpi radialis (wrist extensor), and palmaris longus (wrist flexor). However, we cannot rule out the possibility that recorded EMG signals are influenced by the activity of nearby synergistic muscles. After each implantation, a knot was tied just distal to the electrode exit site to anchor the electrode in place, the needle was removed, and incisions were sutured. Animals were given at least 8 days to recover from the surgery before proceeding to behavior training.

EMG recordings acquired during behavior were amplified and bandpass filtered (1-75,000 Hz) using a 16-channel bipolar amplifying headstage (C3313, Intan technologies). Data was digitized and acquired at 4 kHz using RHD2000 USB interface board and RHD USB interface software (INTAN technologies).

### Optogenetic Inactivation

Optogenetic inactivation was achieved using the same methods described by Koh, Ma *et al.*^45^. After a VGAT-ChR2-eYFP^46^ mouse completed a few climbing sessions with randomly-positioned handholds, it underwent a surgery in which dental cement was removed and a 2 mm craniotomy was made above the left RFA (centered 2.25 mm rostral and 1.00 mm lateral to bregma) or M1 portion of CFA (centered 0.25 mm rostral and 1.5mm lateral to bregma). In experiments without neural recording, a cranial window was implanted over the craniotomy by applying a thin layer of Kwik-Sil (WPI) over the dura followed by a 3 mm diameter #1 thickness cover glass (Warner Instruments). Dental cement sealed the circumference of the glass and protected the remainder of the exposed skull. In some mice, a custom stainless steel ferrule guide (Ziggy’s Tubes and Wires) was cemented above the skull a set distance above the surface of the brain. This distance was calibrated to ensure that the cone of light emanating from a 400 μm core, 0.50 NA optical patch cable terminating in a 2.5 mm ceramic ferrule (M128L01, Thorlabs) would illuminate a spot of light 2 mm in diameter onto the surface of the brain.

Animals typically returned to climbing the next day, however mice sensitive to anesthesia were given an extra day to recover from surgery. During experimental behavior sessions, the optical patch cable was inserted into the ferrule guide, if present, or attached to the arm of a 3-axis micromanipulator (M3-LS-3.4-15-XYZ, New Scale Technologies) to position the fiber tip over the inactivation site. To silence firing throughout cortical layers, we used a 473 nm laser (MDL-III-450-200mW, Opto Engine LLC) to apply 25 ms pulses of blue light at an intensity of 10 mw/mm^2^ in a 2 mm circle on the brain surface^46^. During each session with motor cortex inactivation, light was applied during a random subset of the trials after the bout distance of the wheel exceeded a random threshold between 0॰ and 20॰, and the mouse was actively climbing (informed by wheel encoder data). Inactivation trials were never triggered less than 5 s apart. During the remaining trials, no light was delivered and the trial time was saved to designate a control trial. Our method for randomly triggering laser and control pulses ensured that both types of trials were broadly distributed across the muscle activity states that comprise climbing behavior.

The ratio of laser to control trials, and the total number of each trial type, varied by experiment. For experiments testing the effect of optogenetic inactivation on muscle activity, a 1:2 ratio of laser to control trials was used, and for each mouse, 616-2009 laser and 1280-4047 control trials occurred across 11-20 behavior sessions. Experiments testing the effect of optogenetic inactivation on neural activity used a 1:1 ratio of control to laser trials, and for each mouse, 65-613 laser (median: 253) and 140-656 (median: 256) control trials across 2-4 behavioral sessions each.

### Neural Recording

For experiments with neural recording in one area and optogenetic inactivation in the other, an acute recording using one Neuropixel 1.0 probe (Imec) was performed during daily climbing sessions (2-4 total per animal). Data for experiments with neural recording in both RFA and CFA with no optogenetic inactivation was collected as part of a larger data collection effort in which other brain areas were recorded alongside RFA and CFA. The tip of each probe was sharpened 20॰ prior to use.

To expose each recording area, dental cement was removed from the skull and a 0.5 mm diameter craniotomy was made over RFA and/or CFA. Recording craniotomies were centered 2 mm rostral and -1.25 mm lateral for RFA and 0.5 mm rostral, 1.5 mm lateral for CFA. A Pt/Ir reference wire was implanted to a depth of 1.5 mm in left posterior parietal cortex. Dental cement was carefully applied to cover all remaining exposed skull. After the surgery and between recording sessions, craniotomies were sealed with opaque silicone elastomer (Kwik-Cast, World Precision Instruments). The animal was given 1-2 mL of extra water and allowed to recover overnight or one additional day if necessary (due to anesthesia sensitivity) after the craniotomy procedure.

Prior to each recording session, the animal was head-fixed on the wheel, the silicone elastomer was removed, and each craniotomy was moistened with saline. Each Neuropixel was implanted using a 3-axis multi-probe micromanipulator (see above). Once the probe was fully inserted, the brain surface and exposed skull were covered by a small amount of liquid paraffin oil to maintain hydration of the underlying tissue. Data was acquired at 30 kHz and bandpass filtered at 0.3 to 10 kHz using SpikeGLX software (Janelia Research Campus). Each recording lasted approximately one hour.

### Quantification and Statistical Analysis

All analysis was completed in MATLAB versions 2023b or later (MathWorks). Significance was assessed with alpha = 0.05.

### EMG Processing and Analysis

Raw EMG measurements were high-pass filtered at 250 Hz, rectified, smoothed, downsampled to 1 kHz, and z-scored across time. For experiments with optogenetic inactivation, smoothing was done with a filter where one side of 0 was defined by a Gaussian with a 10 ms standard deviation and the other was set to all zeros. This enabled more precise detection of perturbation onset. For experiments without optogenetic inactivation, a standard Gaussian filter with a 10 ms standard deviation was used.

To compute EMG trial averages, time series segments from -20 to 100 ms around trial onset were collected. Outliers were removed by calculating the total Euclidean distance between each trial segment -20 to 0 ms before trial onset (MATLAB function ‘pdist2’). The mean distance for each trial was computed and a threshold was set one standard deviation above the mean; trials with a mean distance from other trials above this threshold were excluded from subsequent analyses.

EMG trial averages were calculated after subtracting the mean EMG in the baseline period, -20 to 0 ms before trial onset, from each trial. The absolute difference between control and inactivation trials was calculated by taking the difference between the baseline-subtracted means for each muscle. This value was corrected for the difference expected by chance using the following method. On 10000 separate iterations, the absolute difference was calculated between two randomly selected halves of the control trials. The mean absolute difference across the 10000 iterations was then computed and subtracted from the absolute difference time series computed between laser and control trials.

### Identifying EMG trials with high similarity to reaching

Trials were considered “reach-like” if the baseline muscle activity was well-correlated with the average muscle activity around reaching onset. For one animal performing the reaching behavior, 200 ms vectors of muscle activity were assembled by concatenating the 50 ms of trial-averaged normalized EMG from each of the four muscles. These vectors were assembled for 4 overlapping 50 ms segments beginning 15, 10, 5, and 0 ms prior to reach onset to each of the 4 water spouts in the reaching paradigm – resulting in 16 reaching baseline vectors per reaching animal. Then, climbing trials were identified and for each, a 200 ms baseline muscle activity vector was assembled the same was as in reaching using the 50 ms preceding effect onset (with the effect setting in around 10 ms, so -40:10 around light onset). Then, Pearson correlation was calculated between each trial’s baseline muscle activity vector and each of the 16 reaching baseline vectors. If any of these 16 correlation values exceeded a threshold (0.5), the trial was considered “reach-like.”

### Behavior Segmentation

The time of climbing onset and offset was estimated using wheel encoder data. The encoder signal was smoothed with a 1 s Savitsky-Golay filter and converted to angular velocity. A threshold of 0.5°/s was set to detect periods of immobility and wheel turning from the angular velocity. Wheel turning segments shorter than 16° were discarded. The onset was defined as the first instance the wheel turning distance exceeded the 16° threshold for a climbing bout and the offset was defined as the last instance the wheel was turning (ie, above the velocity threshold) before at least 0.3 seconds of immobility (defined as below the velocity threshold).

The onset time of each climbing bout detected in this way was then used to identify the onset of right forelimb muscle activity prior to each bout. For estimating the neural activity onset in relation to muscle onset (Fig. 5f-j), we wanted to use trials in which the animal transitioned from no right forelimb activity to using their right forelimb to turn the wheel within 500 ms. Therefore, we sought to exclude trials that did not fit this criteria, such as those initiated by left forelimb or hindlimb movement. To estimate muscle activity onset prior to all wheel turning onsets, the EMG in the time window -1000 to 1000 ms around each wheel onset was summed across muscles and the sum was smoothed with a standard Gaussian having a 20 ms standard deviation. The change point (Matlab function ‘findchangepts,’ statistic = ‘rms’) of the summed EMG was detected within the time window 500 ms before wheel onset to 200 ms after wheel onset. Trials were then excluded if the mean summed EMG in the time window from 1000 ms prior to wheel onset to the change point exceeded a low threshold (indicating forelimb movement prior to the detected muscle activity onset) or if the change point occurred after the wheel onset (indicating that other limbs drove the initial turning of the wheel).

### Spike Sorting and Unit Curation

Raw neuropixel data collected on SpikeGLX was spike-sorted with Kilosort 3^110^ using default parameters. During some sessions with optogenetic inactivation and neural recording, electrical artifacts were recorded at light onset and/or offset. The light artifact was reduced by excluding channels located outside of the brain from sorting. Occasionally, artifacts were incorrectly identified as spikes and assigned to units; these units were excluded from analysis.

In some mice in which both RFA and CFA were recorded, additional histological verification of each implanted probe location was performed (n = 8 mice, 13 sessions). Each probe was coated with a fluorescent red lipophilic dye CM-DiI (Invitrogen) before each acute recording, leaving a fluorescent red probe tract in tissue that was imaged in serial coronal sections after recording. In addition, recombinant Cholera Toxin Subunit B (CTB) conjugated to Alexa Fluor 488 dye (Invitrogen) was injected into the cervical spinal cord segments C4-C8 to retrogradely infect and label the cell bodies of corticospinal neurons in RFA and CFA after the last recording session, 3-5 days before perfusion. Slices were mounted with a mounting medium containing DAPI (4′,6-diamidino-2-phenylindole) to label cell nuclei (00-4959-52, Invitrogen). Images of coronal brain sections containing each probe tract were registered to the Allen Mouse Brain Common Coordinate Framework (CCFv3)^111^ using the open-source Python package HERBS (Histological E-data Registration in rodent Brain Spaces)^112^ to identify which sections of each probe were in which brain region. Each recorded unit was associated with a waveform centroid positioned along the probe, which was used to localize units to specific brain regions within the CCF. Units assigned to primary motor cortex or secondary motor cortex in sessions where the probe tract was colocalized with corticospinal neuron cell bodies were considered CFA units and RFA units, respectively. In mice in which corticospinal back-labeling was done to define CFA and RFA, probe tracks fell within back-labeled volumes 100% of the time.

Depths of the centroids of recorded units on the probe were used to identify cortical cells in experiments where histology was not performed (n = 6 mice, 13 sessions). For each recording, the depth of the most superficial unit was used as an approximation of the distance from the cortical surface and the deepest electrode on the probe. We excluded any units beyond 1500 μm from the pia based on previous estimates of the thickness of CFA^20^.

Wide-waveform, putative pyramidal cells and narrow-waveform, putative inhibitory interneurons were defined based on waveform width. Waveform widths for each unit were calculated by finding the trough-to-peak duration of the waveform template assigned by Kilosort. Past reports^35,47^ yielded a threshold of 0.417 ms to distinguish between wide-waveform and narrow-waveform units, so we used this threshold to separate the two types. While it is likely that some inhibitory interneurons had wider waveforms than this threshold, previous results using optical tagging to identify GABAergic neurons suggest that the fraction of wide-waveform neurons that are GABAergic is very low^35,113^.

Units of each cell type were further classified by cortical layer. Conservative depth boundaries were established based on the range of depths of units assigned to specific layers of RFA and CFA during CCF registration (Supplementary Fig. 8). For RFA (CFA inactivation), the L2/3 depth boundaries were 100 and 350 µm from pia; L5 boundaries were 450 and 750 µm from pia, and L6 boundaries were 1000 µm and 1400 µm from pia. For CFA, the L2/3 boundaries were 100 and 550 µm from pia; L5 boundaries were 650 and 900 µm from pia, and L6 boundaries were 1100 and 1500 µm from pia.

### Optogenetic Inactivation Effects on Firing Patterns

For all recorded units, instantaneous firing rates were calculated from spike times by smoothing each spike train with a 10 ms standard deviation Gaussian. To compute trial averages, trials were defined by the firing rate time series from each unit -20 to 100 ms around all laser and control events. Outlier trials were removed from each session by calculating the mean Euclidean distance between the neural activity series segments -20 to 0 ms before trial onset. Outlier trials were defined as trials with a mean pairwise distance greater than 3 median absolute deviations from the median (Matlab function ‘rmoutliers’).

To restrict analysis to cells typically active during the behavior, units were selected that had a mean firing rate 35 ms after all control trial onsets greater than 0.5 Hz. In Fig. 2, the animal with the least number of units that met this criteria was used to set the number of units used per animal within each inactivation condition. That number of units was then randomly chosen from the group of eligible units from each other animal. For cell-type and lamina-specific analyses, we wanted to maximize the sample size of groups with limited numbers of units, so we did not use the same number of units per animal. Instead, we used all units that met the above firing rate criterion.

The fractional firing rate change between inactivation and control trials was calculated for individual units and averaged across animals. The fractional change between inactivation and control trials was calculated for each unit by taking the mean control and inactivation trial average across the time window 30-40 ms after trial onset, subtracting the control mean from the inactivation mean, and dividing the resulting value by the control mean. We used non-parametric statistical tests to compare fractional change between groups of cells because distributions of effects did not meet the required assumptions of parametric tests.

To verify that Channelrhodopsin-2 (ChR2) stimulation of inhibitory interneurons only occurred in the region under the light and did not spread to the recorded region, we used a modified version of the Stimulus Associated Latency Test (SALT)^48^. SALT was developed to detect light-responsive neurons in a statistically-based, unsupervised manner; thus negative results from this test indicate a lack of light-responsive neurons in the recorded region. The method was modified slightly to yield a uniform distribution from 0 to 1 under the null hypothesis of no direct light effect as previously described^20^. Narrow-waveform units with average firing rates above 2 Hz across the whole session were selected for each animal and their spike trains 300 ms before to 5 ms after light onset were binned into 5 ms bins around each light event. A p-value was computed for the firing of each unit following light onset, where a low p-value indicates that the neuron is more likely to directly respond to light. The approximately uniform distributions of p-values we observed within single animals and across animals (Supplementary Fig. 3) demonstrate conformance to the null hypothesis of no direct ChR2 activation in the recorded region.

To identify units whose spikes significantly correlated with activation of at least one muscle, instantaneous firing rates were calculated as previously described. Then, the Pearson correlation was measured between each unit’s instantaneous firing rate time series with each muscle’s recorded EMG time series. A bootstrapping method was used to determine which correlations were significant. For each unit, spike trains were circularly shifted by at least ten seconds and no more than ten seconds less than the total duration of the experiment to eliminate temporal correlation between neural and muscle activity time series, and the Pearson correlation was recomputed. This process was repeated 300 times to obtain a null distribution for correlation expected by chance for each unit and muscle. The observed correlations for each unit were then measured against the null distributions using a two-tailed test. If any p-value corresponding to a muscle was significant (p<0.05), the unit was considered significantly correlated to the activity of the given muscle.

### Estimation of neural activity onset time and variance capture with PCA

For each session, trials were first defined as -350 ms to 250 ms around each EMG onset (see above; 18-68 trials per session, 16 sessions, 10 animals), and then the mean firing rate time series across trials was calculated for each unit. Trial averages for each unit were smoothed with a spline-based filter (‘smoothn,’ Matlab File Exchange, smoothing parameter = 1000) and combined across sessions for animals with more than one session. Principal components analysis (PCA) was applied to resultant 600 ms x N units matrices for each area (Matlab function ‘pca’) and the top 3 PCs were selected. Each PC time series was then converted to the absolute change from baseline by subtracting the mean from -350 to -250 ms from onset, taking the absolute value, and then baseline subtracting again. The results were summed for the 3 PCs. The onset time was estimated by detecting the first time point where the trace fell above a threshold; this threshold was set to 7 standard deviations of the mean across the baseline period for the main figure and analysis. Other thresholds tried (Supplementary Fig. 10) were 4 standard deviations and 10 standard deviations, as well as the point with the maximum rate of change across the whole time series (calculated as the first derivative of the time series).

The pre-movement fraction of variance captured in each area was calculated by dividing the area under the summed PC time series from -150 to 0 ms from muscle activity onset by the total area under the summed PC time series from -150 to 150 ms from onset.

### Delayed Latents Across Groups (DLAG)

The application of DLAG^62^ used firing patterns during climbing bouts greater than 3 seconds long, each defined as the time period between wheel rotation onset and offset (see above). Each climbing bout was divided into whole seconds plus some remainder at the end. The first second and last whole second plus the remainder were discarded to avoid analysis of movement initiation and termination. The remaining middle of the climbing bout was divided into non-overlapping second-long epochs (trials). Within each trial, firing rate matrices were assembled for each region by binning each unit’s firing rate in 20 ms windows, resulting in matrices of size N units x 50 time bins. The DLAG model was then fit using these matrices for each trial. The 16 sessions across 10 animals used in Fig. 5 were used for this analysis plus an additional 10 sessions from 4 animals which had simultaneous recordings in RFA and CFA but no histological verification of their location.

For each session, a total of 9 DLAG models were fit by varying the number of across- and within-area dimensions as follows. 5 models were fit with the across-area dimensions held at 4 and the within-area dimensions varying from 2-6, and 4 models were fit with the within-area dimensions held at 4 and across-area dimensions varying from 2-6 (skipping 4 due to redundancy). For each resulting across-area latent variable from each model, bootstrapping was performed by resampling trials 1000 times and refitting the DLAG model with the given delay set to 0 as previously described^62^. Delays were considered significant if less than 5% of bootstrapped samples performed as well as the model with the delay set to its original value, and non-significant delays were excluded from subsequent analysis. Delays were also excluded if they were longer than 60 ms or less than -60 ms, to restrict analysis to delays on the timescale of interactions between RFA and CFA during reaching^20^.

Across-region latents with statistically significant delays were then combined across sessions corresponding to the same animal within each parameter group. Then, within each animal, the fraction of variance captured by RFA-leading and CFA-leading delays in each area was calculated within each session. Results were averaged across sessions for animals with multiple sessions. Finally, the mean variance captured by RFA-leading latents was finally subtracted from the mean variance captured by CFA-leading latents for each animal and then this difference was averaged across animals.

## SUPPLEMENTARY MATERIALS

**Supplementary Figure 1.**
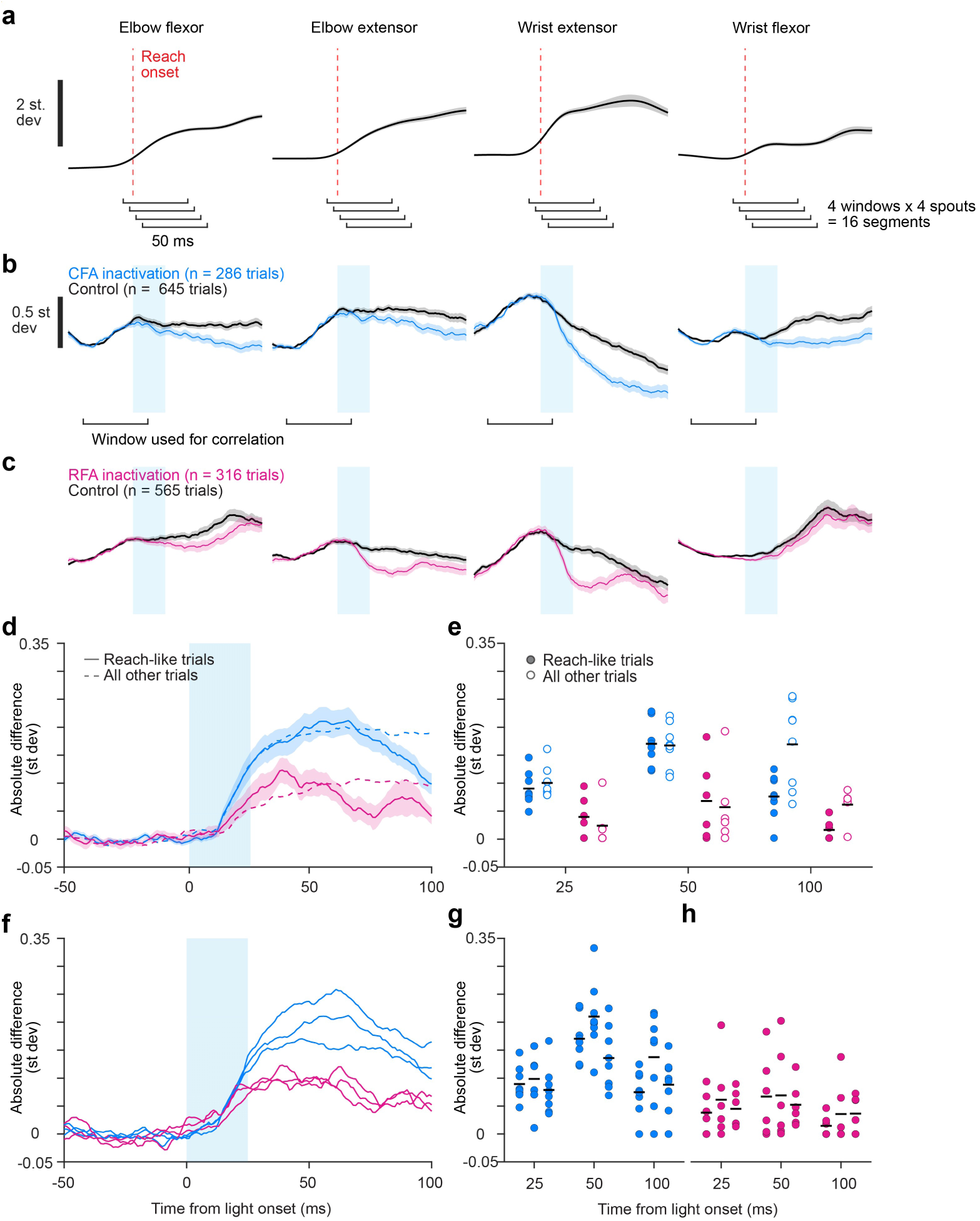
Inactivation effects on climbing trials with reach-like muscle activity. (**a**) Mean +/- SEM normalized muscle activity (units are st dev) for four muscles aligned to reach onset (red dashed line) for one reaching animal. Brackets at the bottom indicate the time segments used for correlation. (**b**) Mean +/- SEM normalized muscle activity for the same four muscles averaged across climbing trials with high correlation to reaching segments (“reach-like”) in (**a**) for control and CFA inactivation in one climbing animal. (**c**) Same as (**b**) but for one animal with RFA inactivation during climbing. (**d**) Mean ± SEM absolute difference between inactivation and control trial averages averaged across all muscles and animals for inactivation (cyan bar) of RFA or CFA during reach-like trials (solid lines). Equivalent means for all other trials are overlaid (dashed). (**e**) Average absolute difference across muscles between inactivation and control trial averages at 25, 50, and 100 ms after trial onset for individual animals (circles) and the mean across animals (bars) for climbing reach-like trials (filled circles) and all other trials (open circles) after inactivation of RFA or CFA. (**f**) Mean absolute difference between inactivation and control trial averages averaged across all muscles and animals for inactivation (cyan bar) of RFA or CFA during reach-like trials, with each trace corresponding to trials correlated to reaching trials of a different reaching animal (3 total). (**g,h**) Average absolute difference across muscles between inactivation and control trial averages at 25, 50, and 100 ms after trial onset for individual animals during reach-like trials correlated with reaching muscle activity of different reaching animals, for CFA inactivation (**g**) and RFA inactivation (**h**).

**Supplementary Figure 2.**
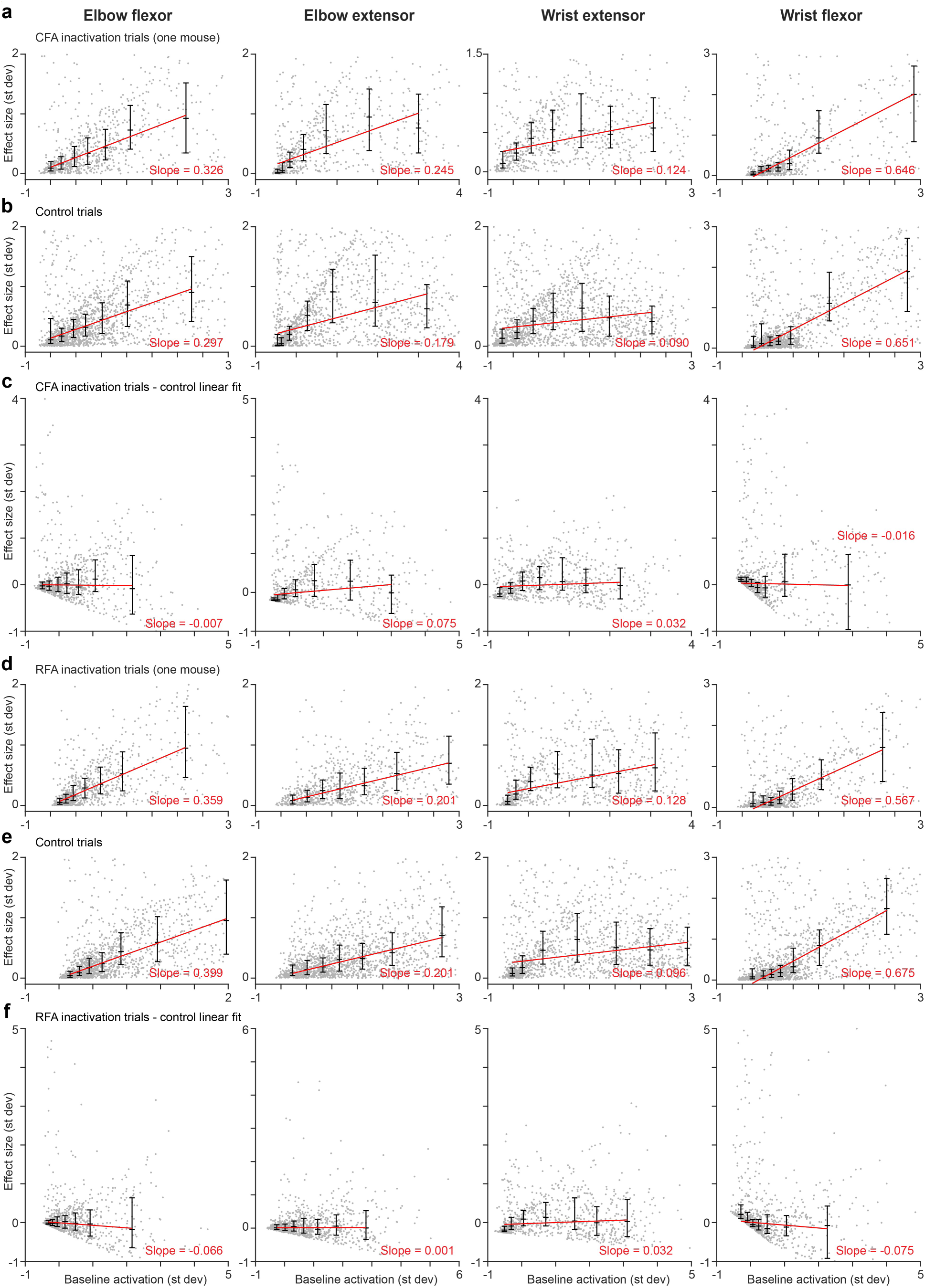
Inactivation effect size versus baseline muscle activation. (**a**) For one animal, scatterplot of the inactivation effect size, defined as the absolute difference between baseline muscle activation (mean -40 to 10 ms around light onset) and muscle activation averaged 20 to 30 ms after trial onset, versus baseline activation for each muscle (columns) across all CFA inactivation trials (n=742). Whisker plots reflect the 1^st^, 2^nd^ and 3^rd^ quartile effect sizes for data points divided into 7 equally sized bins by baseline activation. Each whisker plot is centered at the median baseline activation for data points in each given bin. A linear fit was made to the median effect sizes for the seven bins (red). (**b**) Same as (**a**) but for all control trials (n=1546). (**c**) Same as (**a**) but after subtracting the linear fit to control trials in (**b**) from each inactivation trial. Note that no strong linear trend remains, suggesting that the effect sizes are not strongly determined by baseline muscle activation. (**d-f**) Same as (**a-c**) but for one animal with RFA inactivation (832 inactivation trials, 1506 control trials). Similar results were seen in other animals. Additional analysis probing for correlation between CFA inactivation effect size and baseline muscle activation is provided in Koh, Ma *et al.*^45^. Little was found.

**Supplementary Figure 3.**
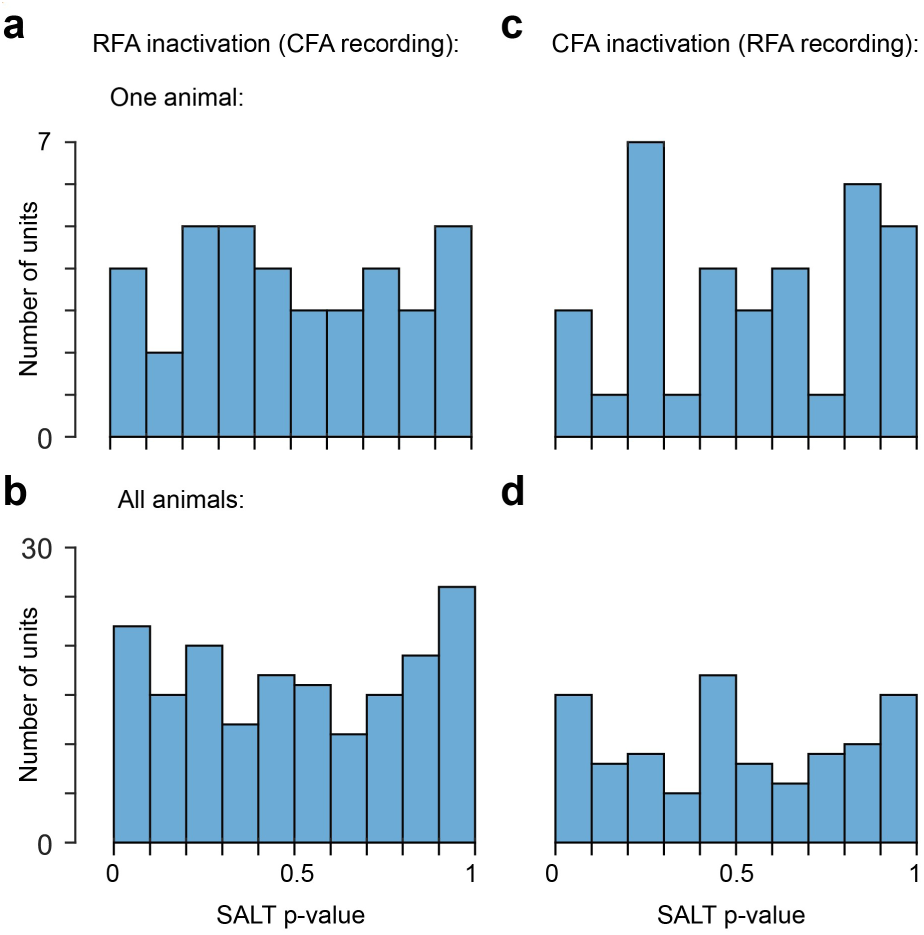
No evidence for off-target light effects during optogenetic experiments. (**a-d**) Histograms of p-values from the modified SALT method for narrow-waveform neurons recorded in all sessions for one animal (**a,c**) or all sessions across all 4 animals (**b,d**) in either CFA after RFA inactivation (**a,b**) or RFA after CFA inactivation (**c,d**). The approximately uniform distributions here affirm the null hypothesis that neurons are not directly activated by light in the off-target region.

**Supplementary Figure 4.**
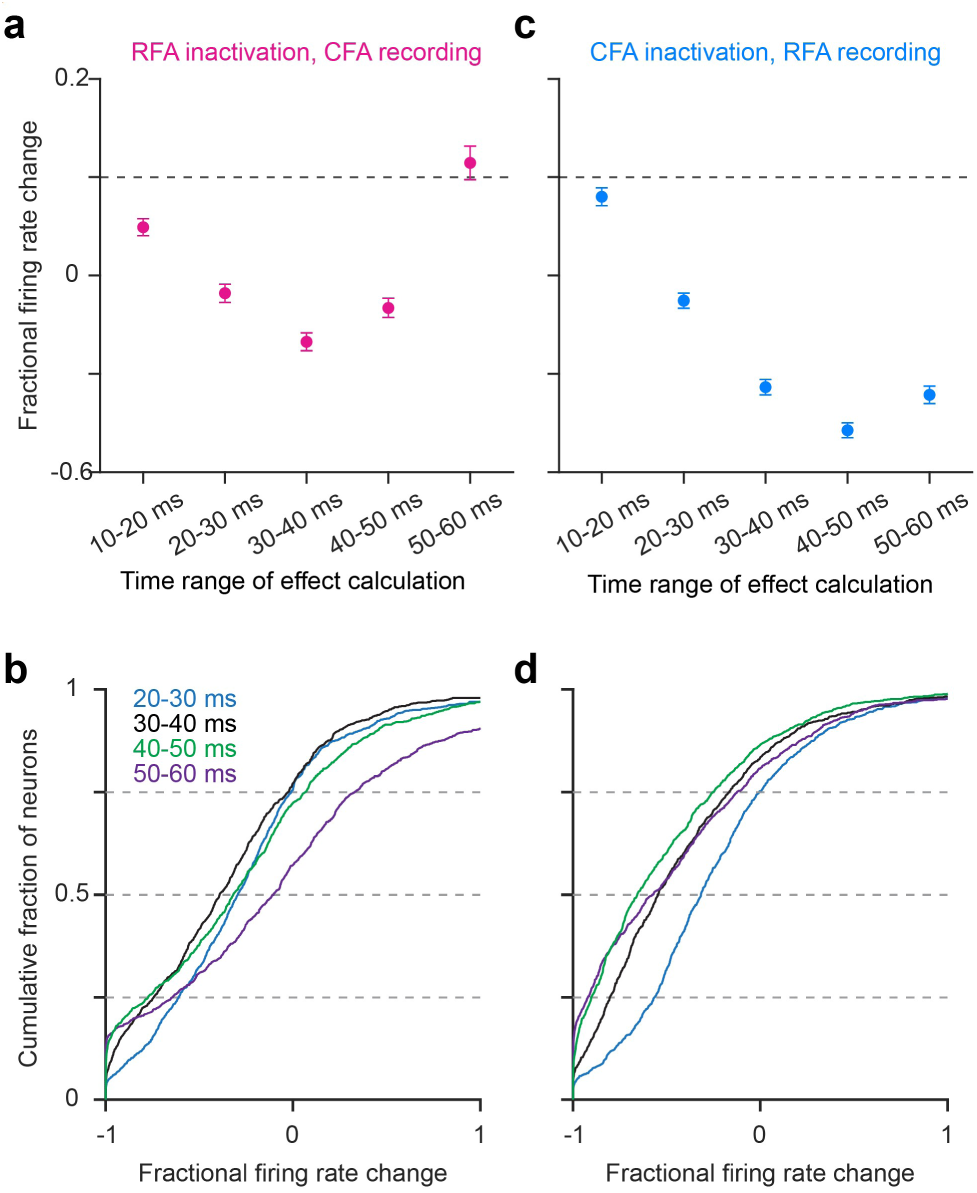
Fractional change calculated in different time windows. (**a,c**) Mean ± SEM fractional change between control and inactivation trials calculated in different time ranges after trial onset across neurons in CFA after RFA inactivation (**a**) or in RFA after CFA inactivation (**c**). (**b,d**) Cumulative histograms of the fractional change in firing rate between inactivation and control trials in 10 second window centered at different times after trial onset for neurons in CFA after RFA inactivation (**b**) or in RFA after CFA inactivation (**d**).

**Supplementary Figure 5.**
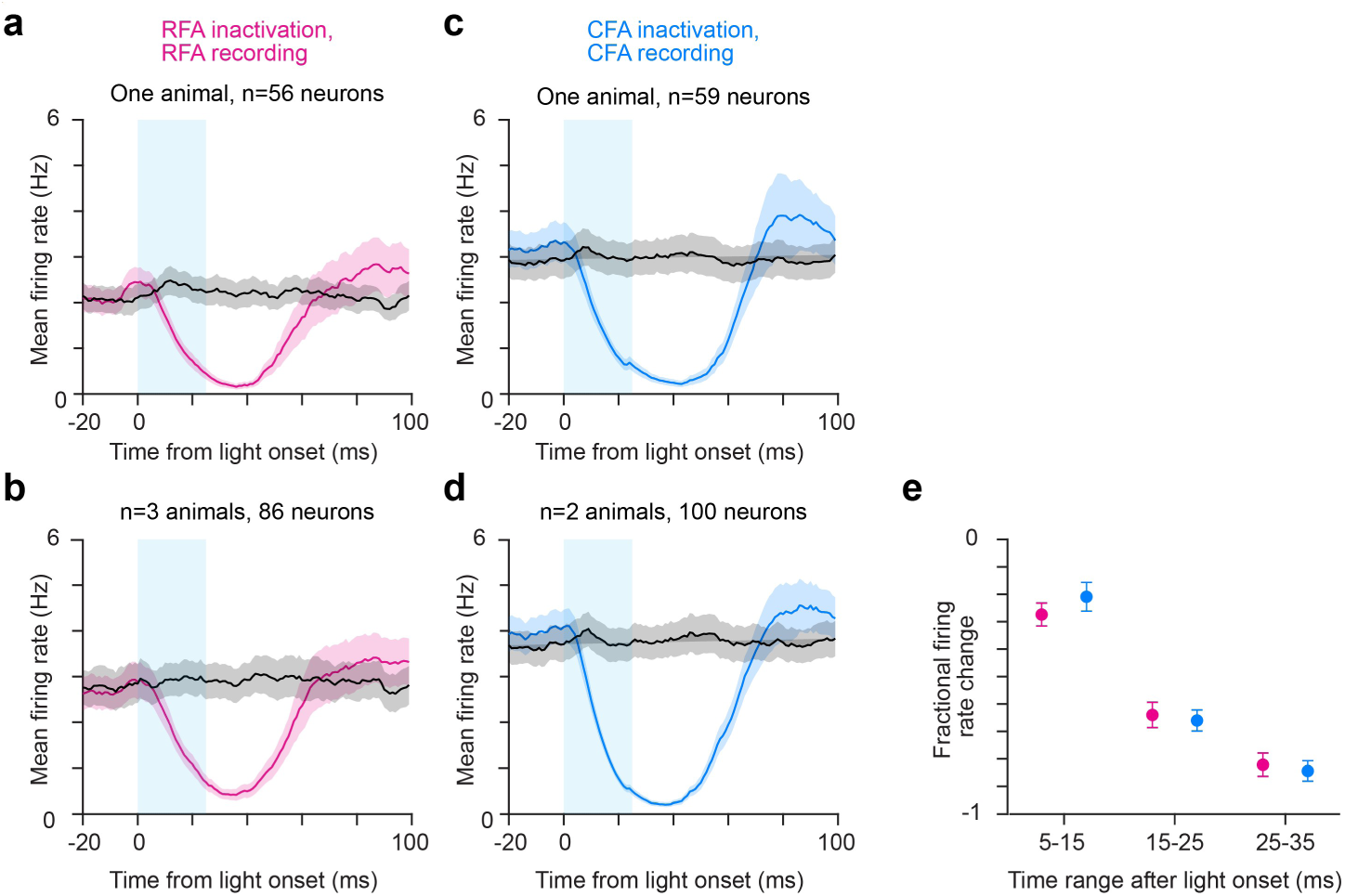
Local inactivation is similarly complete in both areas. (**a**,**b**) Mean ± SEM firing rate for cells from one animal (**a**) after inactivating RFA (cyan bar) and recording in RFA (**a**) or multiple animals (**b**) and for control trials in each. (**c,d**) Mean ± SEM firing rate for cells from one animal (**a**) after inactivating CFA (cyan bar) and recording in CFA (**a**) or multiple animals (**b**) and for control trials in each. (**e**) Mean ± SEM fractional change between control and inactivation trials across neurons in RFA (magenta) and CFA (blue) after local inactivation in different time windows following trial onset.

**Supplementary Figure 6.**
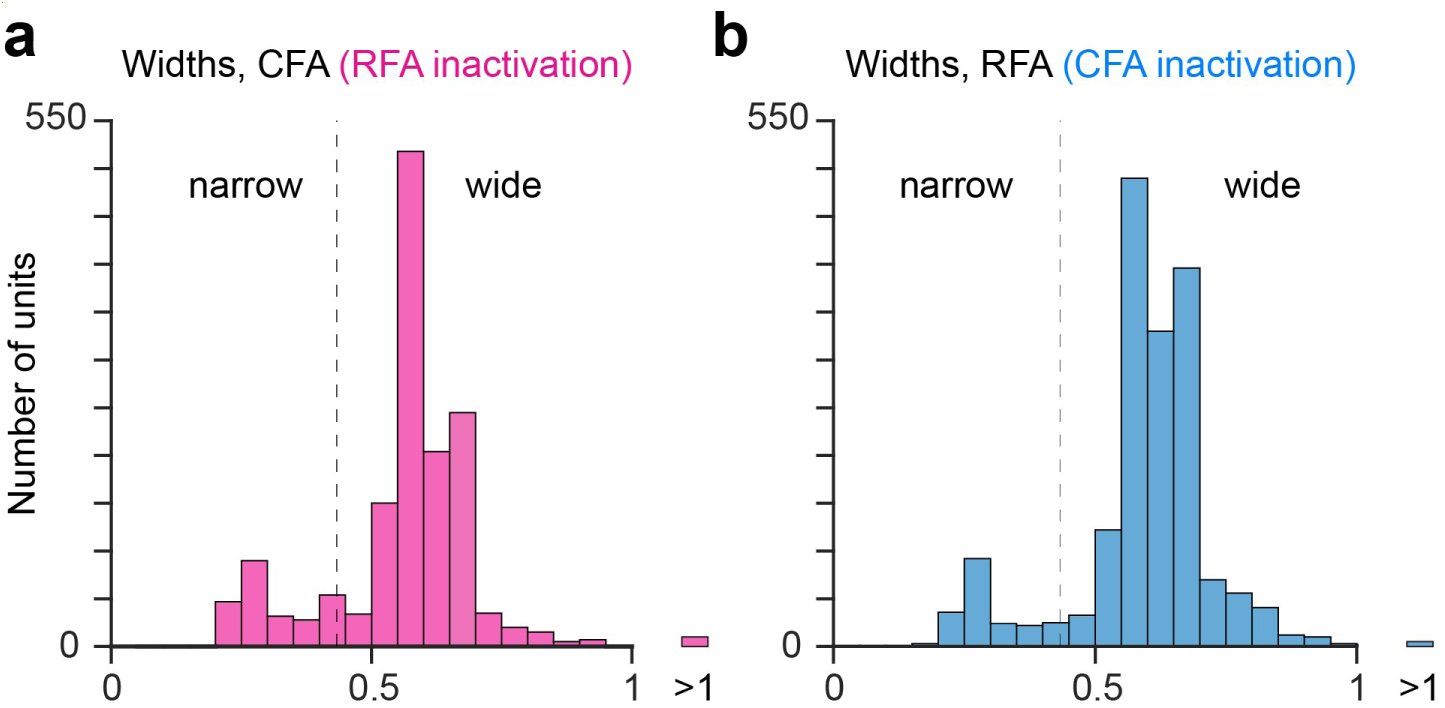
Distribution of waveform widths among recorded units in RFA and CFA. (**a, b**) Histogram of spike waveform widths in CFA (**a**) and RFA (**b**) recorded during optogenetics experiments. Vertical dashed line corresponds to the threshold dividing wide- and narrow-waveform cells in this study.

**Supplementary Figure 7.**
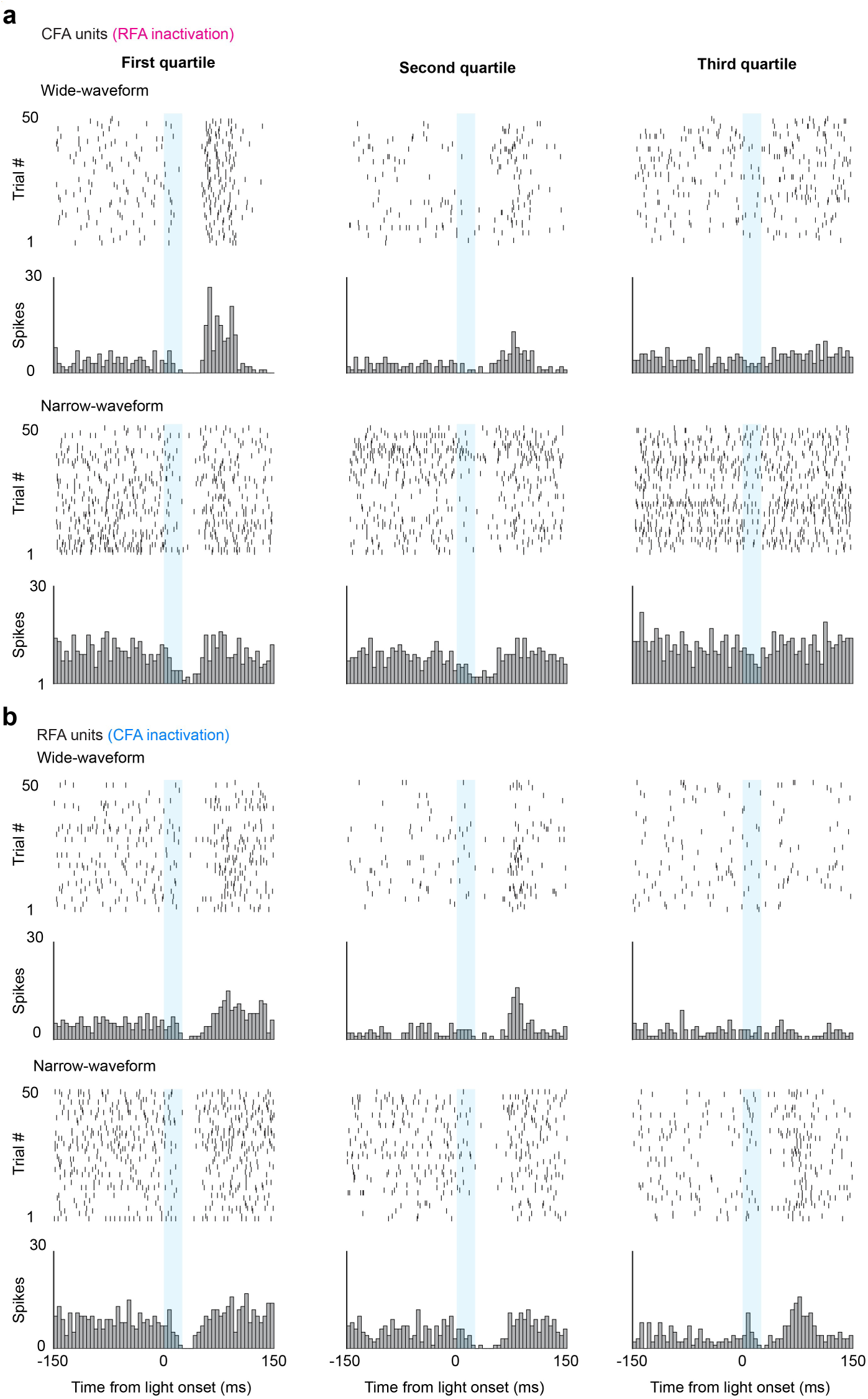
Examples of single wide- and narrow-waveform units recorded during inactivation of the other area. (**a,b**) Spike rasters and peri-stimulus spike histograms of wide- and narrow-waveform CFA units recorded during RFA inactivation (**a**) and RFA units recorded during CFA inactivation (**b**) from the top three quartiles of fractional change effect size, where the first quartile corresponds to the largest effect size.

**Supplementary Figure 8.**
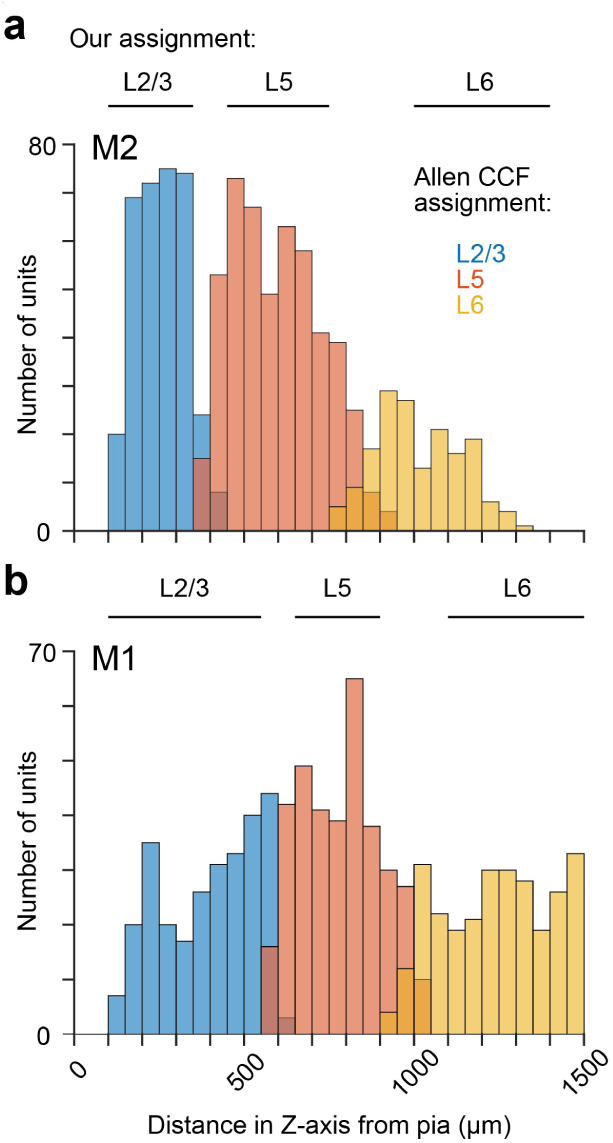
Depth ranges for layer assignments in M2 (RFA) and M1 (CFA). (**a,b**) Histogram of the depth from the pia surface of units assigned to specific cortical layers of M2 (**a**) and M1 (**b**) by the Allen Common Coordinate Framework^111^. Units were combined across 16 recording sessions from 10 animals. Conservative depth boundaries used in this study are shown above each histogram.

**Supplementary Figure 9.**
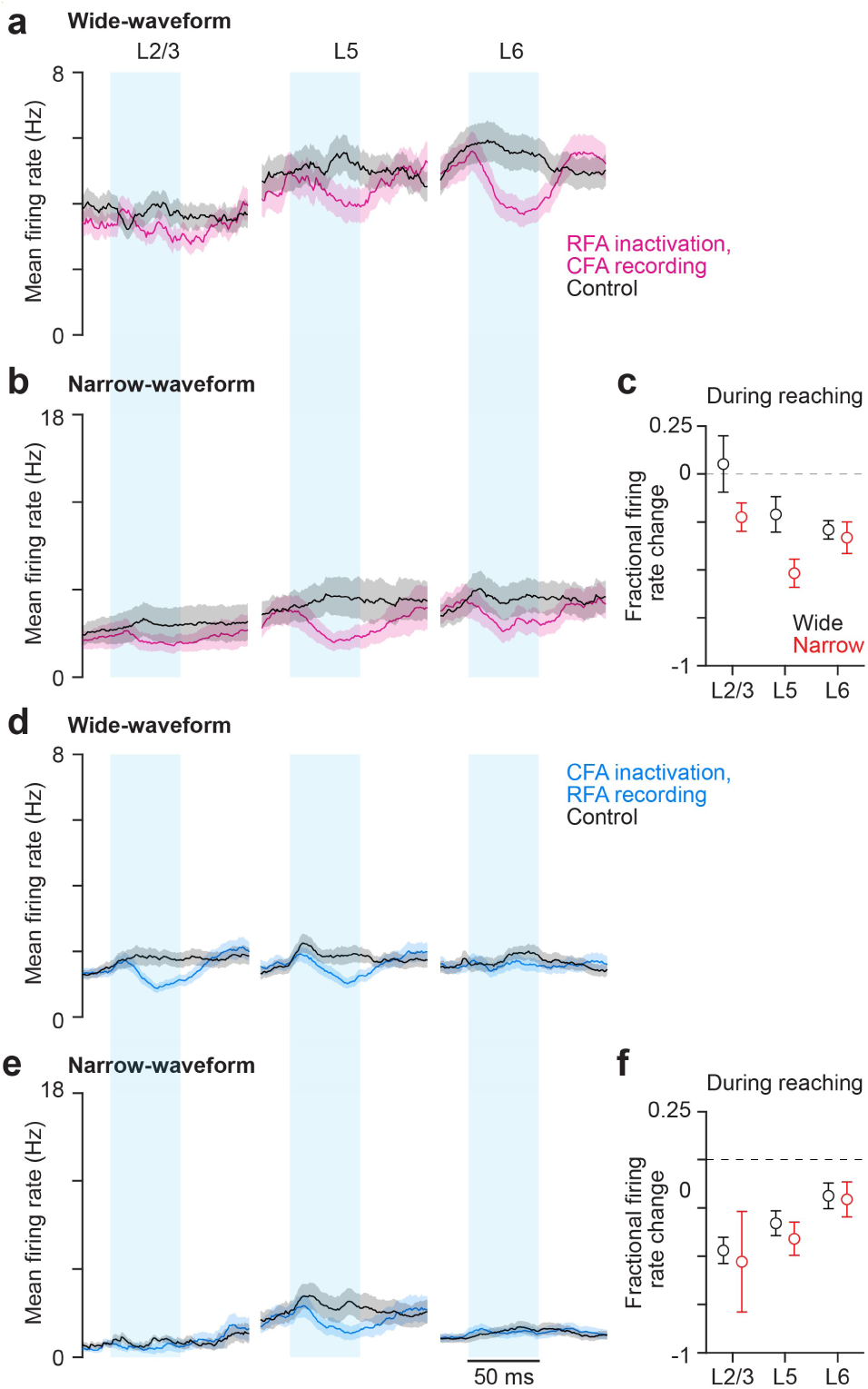
Effects of inactivation across layer groups during reaching. (**a**,**b**) Mean ± SEM firing rate for wide- (**a**; 126 in L2/3, 105 in L5, 240 in L6) L5 (n=105) and narrow-waveform (**b**; 93 in L2/3, 47 in L5, 54 in L6) neurons in CFA assigned to each laminar cell group after RFA inactivation, and for control trials. (**c**) Mean ± SEM fractional firing rate change between inactivation and control trials 35 ms after trial onset for all neurons in different laminar cell groups in CFA after RFA inactivation during reaching. (**d**-**f**) Same as (**a**-**c**), but for RFA neurons (wide: 153 in L2/3, 192 in L5, 143 in L6; narrow: 33 in L2/3, 79 in L5, 55 in L6) recorded during CFA inactivation, instead of vice versa.

**Supplementary Figure 10.**
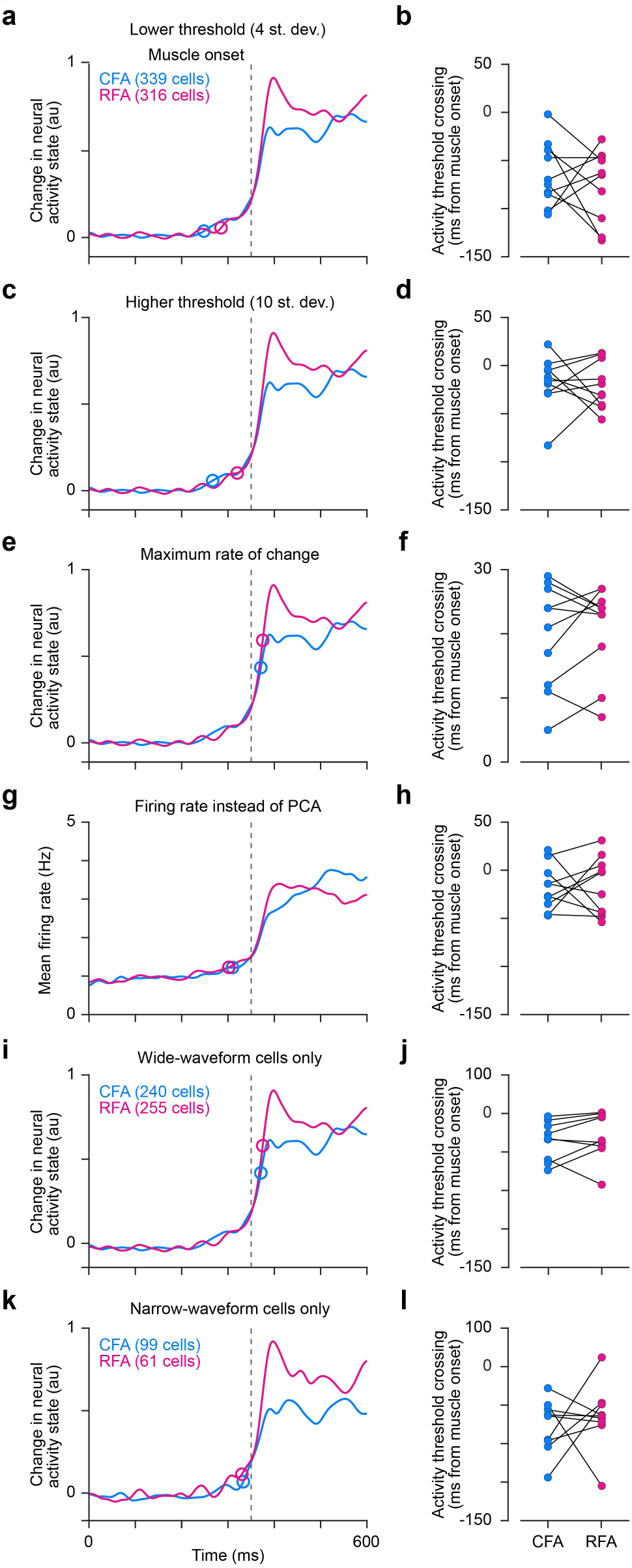
Alternative methods of neural activity onset detection. **(a)** For an example mouse, the normalized activity change from baseline summed across the top three principal components (PCs) for all recorded CFA or RFA neurons, and the top PC for muscle activity surrounding muscle activity onset. Circles indicate the time of estimated activity onset in each area. The threshold for onset detection was 4 st dev of the baseline. **(b)** Time from muscle activity onset at which the neural activity change from baseline exceeded a low threshold for each animal using the onset detection method in (**a**). (**c,d**) Same as (**a,b**) but with a onset detection threshold of 10 st dev of the baseline. (**e,f**) Same as (**a,b**) but with onset detected as the point of the maximum rate of change. (**g,h**) Same as (**a,b**) but with the mean firing rate across neurons used instead of PCA and the onset threshold set at 7 st dev of the baseline. (**i,j**) Same as (**a,b**) using wide-waveform cells only in each area and with an onset detection threshold of 7 st dev of the baseline. (**k,l**) Same as (**i,j**) but for narrow-waveform cells only in each area.

